# Proteomics of spatially identified tissues in whole organs

**DOI:** 10.1101/2021.11.02.466753

**Authors:** Harsharan Singh Bhatia, Andreas-David Brunner, Zhouyi Rong, Hongcheng Mai, Marvin Thielert, Rami Al-Maskari, Johannes Christian Paetzold, Florian Kofler, Mihail Ivilinov Todorov, Mayar Ali, Muge Molbay, Zeynep Ilgin Kolabas, Doris Kaltenecker, Stephan Müller, Stefan F. Lichtenthaler, Bjoern H. Menze, Fabian J. Theis, Matthias Mann, Ali Ertürk

## Abstract

Spatial molecular profiling of complex tissues is essential to investigate cellular function in physiological and pathological states. However, methods for molecular analysis of biological specimens imaged in 3D as a whole are lacking. Here, we present DISCO-MS, a technology combining whole-organ imaging, deep learning-based image analysis, and ultra-high sensitivity mass spectrometry. DISCO-MS yielded qualitative and quantitative proteomics data indistinguishable from uncleared samples in both rodent and human tissues. Using DISCO-MS, we investigated microglia activation locally along axonal tracts after brain injury and revealed known and novel biomarkers. Furthermore, we identified initial individual amyloid-beta plaques in the brains of a young familial Alzheimer’s disease mouse model, characterized the core proteome of these aggregates, and highlighted their compositional heterogeneity. Thus, DISCO-MS enables quantitative, unbiased proteome analysis of target tissues following unbiased imaging of entire organs, providing new diagnostic and therapeutic opportunities for complex diseases, including neurodegeneration.

**Graphical Abstract:** 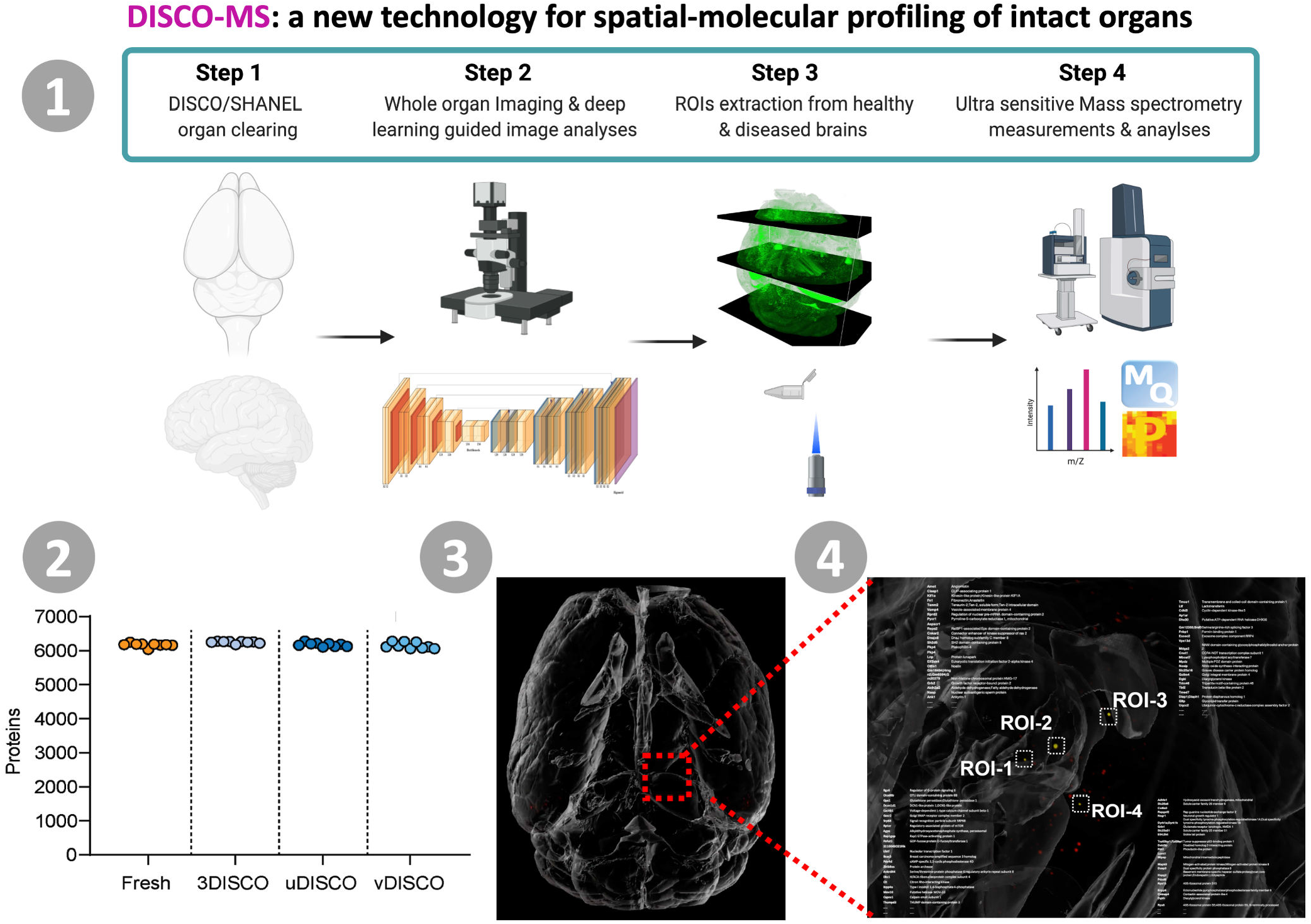

**Highlights:**

1. DISCO-MS combines tissue clearing, whole-organ imaging, deep learning-based image analysis, and ultra-high sensitivity mass spectrometry
2. DISCO-MS yielded qualitative and quantitative proteomics data indistinguishable from fresh tissues
3. DISCO-MS enables identification of rare pathological regions & their subsequent molecular analysis
4. DISCO-MS revealed core proteome of plaques in 6 weeks old Alzheimer‘s disease mouse model Supplementary Video can be seen at: http://discotechnologies.org/DISCO-MS/

## INTRODUCTION

At their early stages, many diseases start with a modest pathology in mostly unknown tissue regions making them hard to identify and characterize. For example, early changes in dementia may include the activation of few local inflammatory cells, changes in the microvasculature, and the appearance of just a few initial amyloid-beta plaques in unknown brain regions (Braak et al., 2011). Such small regional changes are extremely hard to be identified using standard histology, limiting our ability to investigate initial stages of the disease for early diagnostic and therapeutic opportunities. Recent advances in tissue clearing technologies allow fluorescence imaging of complete biological tissues including mouse organs and whole bodies, as well as intact human organs (Belle et al., 2017; Ueda et al., 2020; Zhao et al., 2020). After entire tissues are rendered transparent, end-to-end laser scanning microscopy reveals their cellular and sub-cellular level details. Leveraging artificial intelligence (AI)-guided image analysis, years of imaging work can now be completed within days, allowing to capture even tiny changes in cellular structures, which would be missed in otherwise pre-selected tissue sections (Moen et al., 2019; Sullivan et al., 2018). However, visually pinpointing these regions alone does not answer molecular level, mechanistic questions. Here, we report a new technology in which we apply ultra-high sensitivity MS-based proteome analysis to target tissues comprising less than 100 cells after their identification by organic solvent-based organ clearing and imaging (3-Dimensional Imaging of Solvent Cleared Organs profiled by Mass Spectrometry: DISCO-MS). Using DISCO-MS, we analyzed proteomes of rare pathological regions with known spatial location and morphology in whole mouse brain scans, which provides unique opportunities for spatial-molecular profiling of intact organs (**Movie S1**).

## RESULTS

### MS-based proteomics of solvent-cleared tissue

Tissue clearing enables rapid imaging of intact organs at the cellular level without sectioning. It is a chemical process that relies on tissue permeabilization and subsequent extraction of different biomolecules including water (in organic-solvent-based DISCO clearing methods) and lipids (in most clearing methods) (Ueda et al., 2020). We asked how the proteome of solvent-cleared tissues changes after these diverse and harsh extraction steps. Toward this goal, we turned to MS-based proteomics, which can provide unbiased in-depth insights into the composition, structure, and function of the entirety of expressed proteins (Aebersold and Mann, 2016).

We first explored the feasibility of subjecting solvent-cleared rigid tissue to proteomics sample preparation workflow regarding protein recovery, qualitative and quantitative reproducibility. Using fresh-frozen tissues as controls, we started with 3DISCO and uDISCO tissue clearing methods, two commonly used organic solvent-based transparency methods that are quick and known to provide the highest tissue and organ transparency (Ueda et al., 2020). We tested several protein solubilization approaches based on: Sodiumdeoxycholate (SDC) alone, 2,2,2-Trifluoroethanol (TFE) + Sodiumdodecylsulfate (SDS), and SDS + SDC (See methods for in-depth description). Remarkably, the combination of SDS-based protein solubilization and tissue pulverization, several boiling steps, and acetone-based precipitation/ washing, followed by SDC-resolubilization and protein digestion, yielded qualitatively and quantitatively very similar proteomes among fresh and these two organic solvent-based clearing methods. We identified up to 5,500 proteins across conditions with Pearson correlation coefficients between 0.89 and 0.99. Note that this minimal quantitative variability also reflects biological differences as the cleared brains came from different mice (**Fig. S1**).

While 3DISCO and uDISCO work well for fluorescent dye imaging, the signal of endogenous fluorescence proteins such as EGFP can decay before panoptic imaging (Richardson and Lichtman, 2015). To avoid this, we recently developed vDISCO, which uses fluorescent dye-conjugated nanobodies to stabilize and enhance endogenous fluorescence protein signals (Cai et al., 2019). As vDISCO includes several additional tissue labeling and clearing steps that might change the proteome, we tested our proteomics workflow also on tissues subjected to this technique. Identification of more than 6,000 proteins across all clearing conditions including vDISCO and replicates with Pearson correlations ranging from 0.85 to 0.94 confirmed the feasibility of our workflow (**Fig. 1A**). Next, we asked if we can also analyze archival human brain tissues with the same protocol. To this end, using 3DISCO, uDISCO and SHANEL methods (Zhao et al., 2020), we cleared human brain tissues stored in formalin for more than five years and subjected them to our workflow. Encouragingly, we identified more than 5,000 proteins in all clearing conditions compared to PFA-fixed controls at high quantitative reproducibility (R = 0.91-0.96) (**Fig. 1B**). We concluded that our new sample preparation workflow allowed the MS-based proteome analysis of cleared mouse and human organs at qualitative depth and quantitative accuracy comparable to fresh and PFA-fixed control samples.

**Fig. 1:**
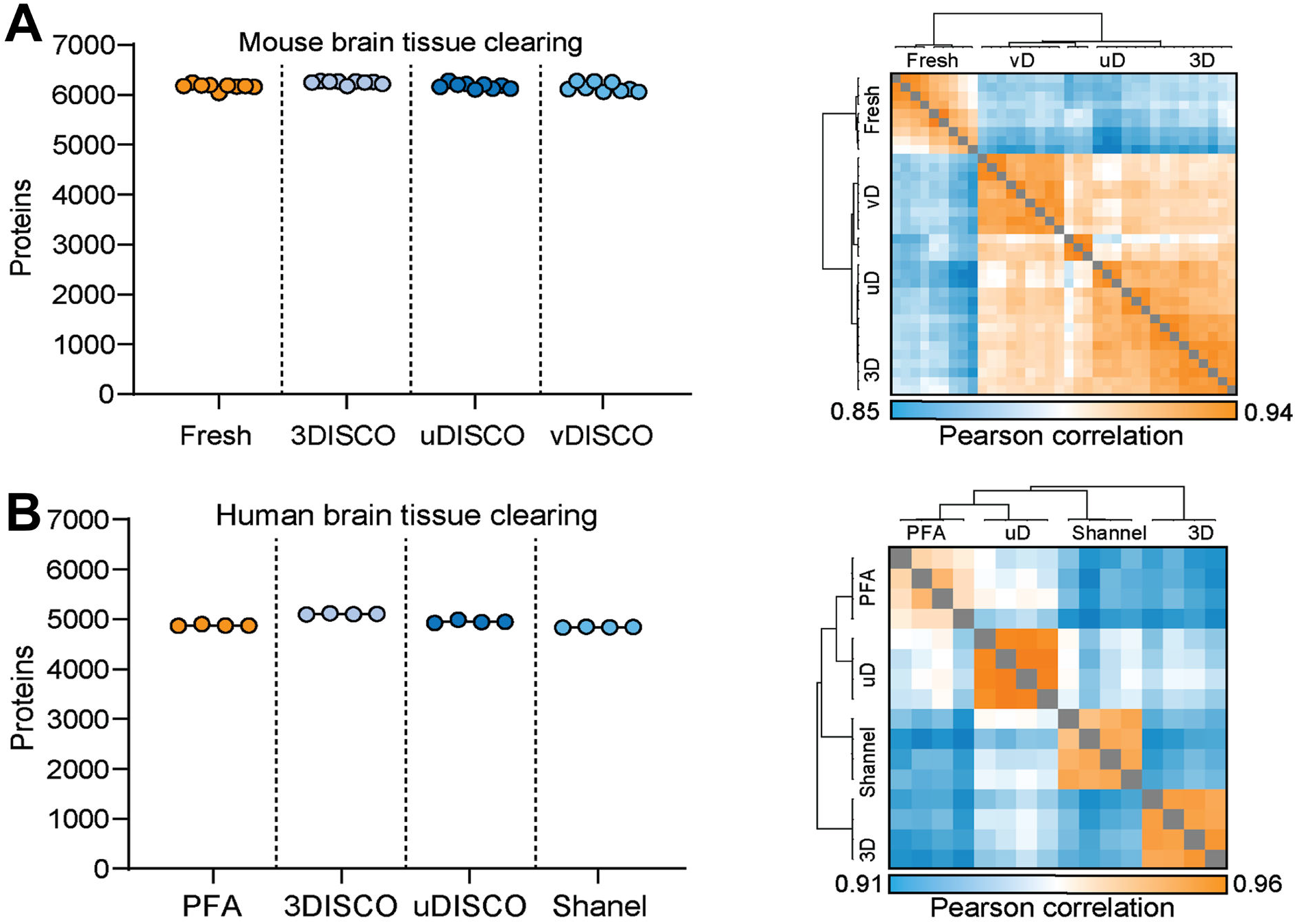
Proteome of cleared rodent and human tissues. **(A)** Proteome analysis from mouse brain tissues after three different organic solvent-based tissue clearing methods (3DISCO, uDISCO, vDISCO) vs. fresh tissue controls. Protein identifications and proteome correlations across all clearing techniques and fresh tissue are shown. (N=3 biological replicates, n=9 total experimental replicates per condition). **(B)** Archived human brain cortex blocks cleared with 3DISCO, uDISCO and SHANEL methods and number of detected protein groups with proteome correlations across all clearing techniques are compared with the detected numbers in PFA fixed blocks. n=4 experimental replicates.

### High proteome yield in vDISCO cleared tissues

Next, we examined the proteome of cleared tissue in-depth to investigate potential protein depletions introduced by the clearing process. Three biological replicates of vDISCO-cleared mouse brains and three fresh-frozen control brains (C57BL/6J) were subjected to our sample preparation workflow for bottom-up proteomics, separated into 16 fractions each and subjected to MS-based proteome analysis and data mining (**Fig. 2A**). This workflow identified close to 8,000 proteins in total across conditions and biological replicates with a high quantitative reproducibility between conditions across the full dynamic range (R = 0.94; **Fig. 2B, Fig. S2**). Furthermore, coefficients of variation within fresh and clearing conditions were below 0.2, highlighting that vDISCO clearing yields proteomes which are qualitatively and quantitatively in the same range as fresh tissue and very reproducible across biological replicates (**Fig. S2**). The only altered gene ontology (GO) term was ‘Blood microparticle’ proteins, which was expected as vDISCO tissues were perfused, while other GO keywords, such as ‘Aging’, ‘Neurogenesis’, ‘Immunity’, ‘Wound healing’, ‘Virus-Host’, ‘Neurodegeneration’, and ‘Receptor’ are quantitatively and qualitatively preserved (**Fig. 2C, Fig. S3**). Cellular proteomes can be quantified by the proteome ruler algorithm, which uses the fixed ratio of total histones to the genome (Wiśniewski et al., 2014) allowing us to relate gene ontology annotations of subcellular localization to protein mass distributions between fresh and vDISCO mouse brains for membrane, organelle, and cytoskeleton terms. We found that all percentage protein mass differences were well below 15% across sub-terms and the protein mass change associated with membrane-related terms was below 3% (**Fig. 2D**). In summary, even the strongest organic solvent-based tissue clearing approach, vDISCO, yields very similar proteomes compared to the fresh tissue.

**Fig. 2:**
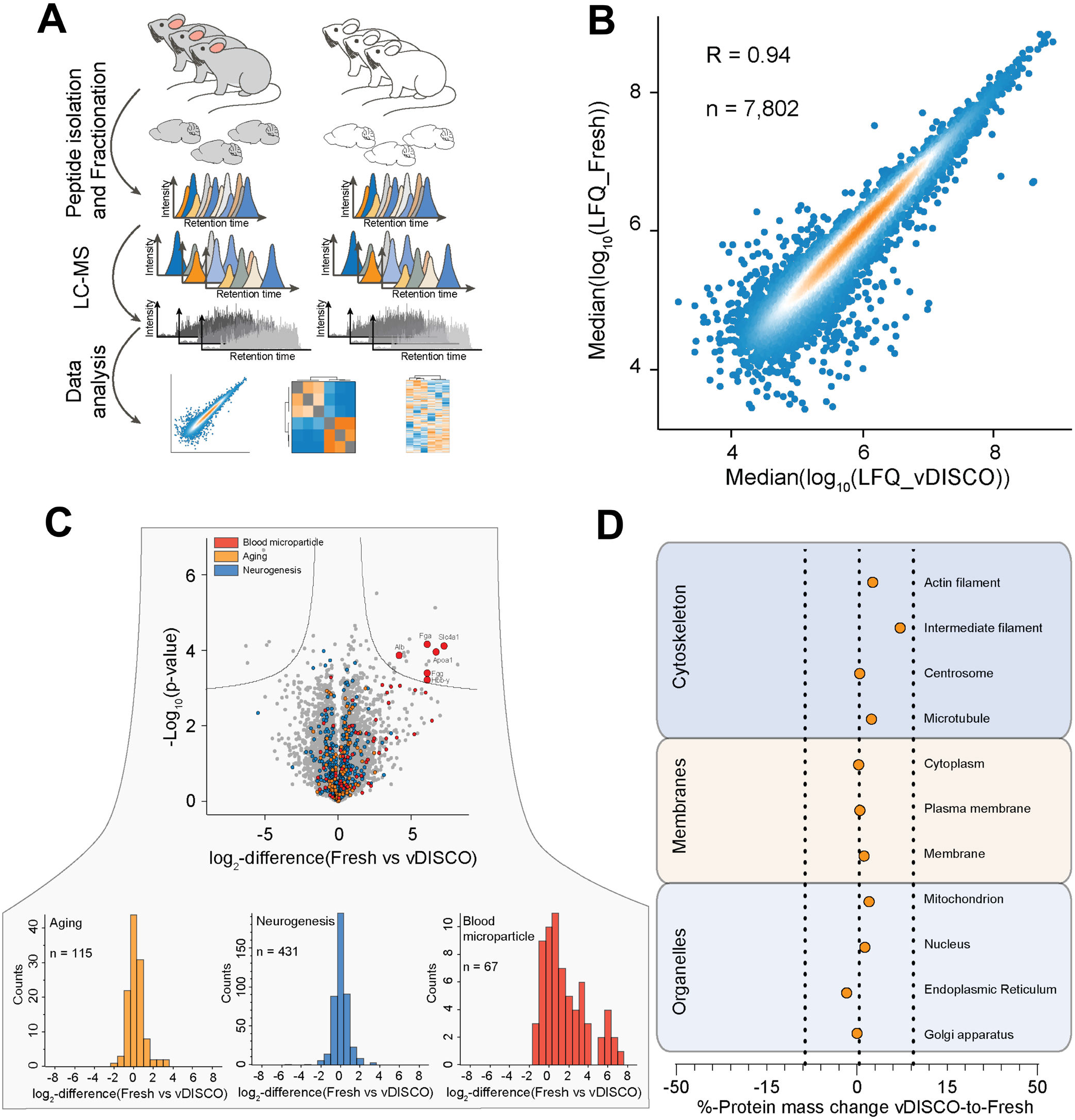
Deep proteome analysis after vDISCO tissue clearing. **(A)** Experimental design for deep proteome analysis after vDISCO clearing vs. snap-frozen fresh tissues. **(B)** Quantitative reproducibility of vDISCO-cleared vs. fresh samples. **(C)** Differential expression analysis of vDISCO-cleared vs. fresh sample proteomes highlighting the expected change in ‘blood microparticle’ due to blood-flushing step for tissue clearing in contrast to fresh samples with retained-blood. Otherwise, proteins in other gene ontology groups were unchanged (See also **Fig. S3**). **(D)** Percentage change of protein mass distribution between vDISCO-cleared vs. fresh biological replicates. Percentage changes are shown as a median change within one group for organelles, membranes and cytoskeleton gene ontology terms (N=3 biological replicates, n=9 total experimental replicates per condition).

### Proteomes of micro-dissected tissues imaged in 3D

After establishing a high-quality and reproducible MS-compatible sample preparation workflow for solvent-cleared tissues, we turned to the proteome analysis of tiny target tissue regions previously imaged and located in 3D. Here, we encountered two major challenges: 1) reliable dissection of small tissue regions identified by 3D-imaging of cleared tissues, and 2) measuring a deep proteome from only a few nano-grams of dissected and rigid solvent-cleared tissue. To solve the first problem, we developed a series of steps to render cleared rigid tissue soft for precise cryosectioning and laser capture microdissection (LCM) without deformation. In short, we reversed the clearing protocol, rehydrated the cleared tissue stepwise, and quickly cryo-preserved it using isopentane in a sucrose bed. This workflow avoided rupturing of the tissue during cryosectioning and allowed us to laser micro-dissect tissue regions as small as 0.0005 mm^3^ corresponding to approximately 60 cells in volume. Next, we miniaturized our sample preparation workflow and then performed MS-based proteomics analysis on a modified trapped ion mobility mass spectrometry platform developed to the highest sensitivity as we recently described for the analysis of single FACS sorted cells (Brunner et al., 2021).

To explore the potential of our technology in clinically relevant applications, we first used a mild traumatic brain injury (mTBI) mouse model to identify and analyze proteomes of brain regions depicting discrete local inflammation. mTBI, including concussions, are common and can lead to long-term comorbidities such as sleep disorders, neuropsychiatric disorders, and even early onset of dementia (Langlois et al., 2005). They are characterized by chronic inflammation, which can induce neurodegeneration in selected brain regions, particularly along the stretched axonal tract (Ertürk et al., 2016). We used a repetitive mTBI injury model on CX3CR1-EGFP mice (**Fig. 4SA–D**), in which all microglia are labelled with an EGFP-fusion construct. The ClearMap quantification approach (Renier et al., 2016) readily identified activated microglia with enlarged morphology in diverse brain regions, especially along the axonal tracts including the optic tract and the corpus callosum (**Fig. S4E–J**). Applying the same mTBI injury model on Thy1-GFP-M reporter mice (expressing GFP only in neurons), we confirmed the axonal abnormalities in the same brain regions (**Fig. 4SK–N**). We then used DISCO-MS workflow on isolated regions of interest (ROIs) including locally activated microglia with known spatial information (**Fig. 3A,B**). Analyzing three ROIs from the optic tract as small as 0.0005 mm^3^ compared to corresponding regions in sham control animals, we quantified up to 1400 proteins per ROI. Principal Component Analysis (PCA) separated the proteomes of ROIs between mTBI and controls (**Fig. 3C**). Several proteins related to axonal damage and repair were upregulated in the mTBI ROIs including Stmn1 (32-fold increase) and Ncan (30-fold increase) (**Fig. 3D**). We found 602 common proteins in all ROIs of mTBI and sham. Comparing ROIs from mTBI among themselves, we found a common proteome signature comprising of a total of 717 proteins (**Fig. 3E**; all the data showing common and unique proteins at each ROI are available at PRIDE repository, see methods for details). We further validated the enrichment of Stmn1 and Ncan in mTBI brain tissues by immunofluorescence (**Fig. 3F-H**). Our data demonstrate that DISCO-MS is a powerful approach to obtain unbiased proteomics information on heterogeneous tissue regions with known spatial location.

**Fig. 3:**
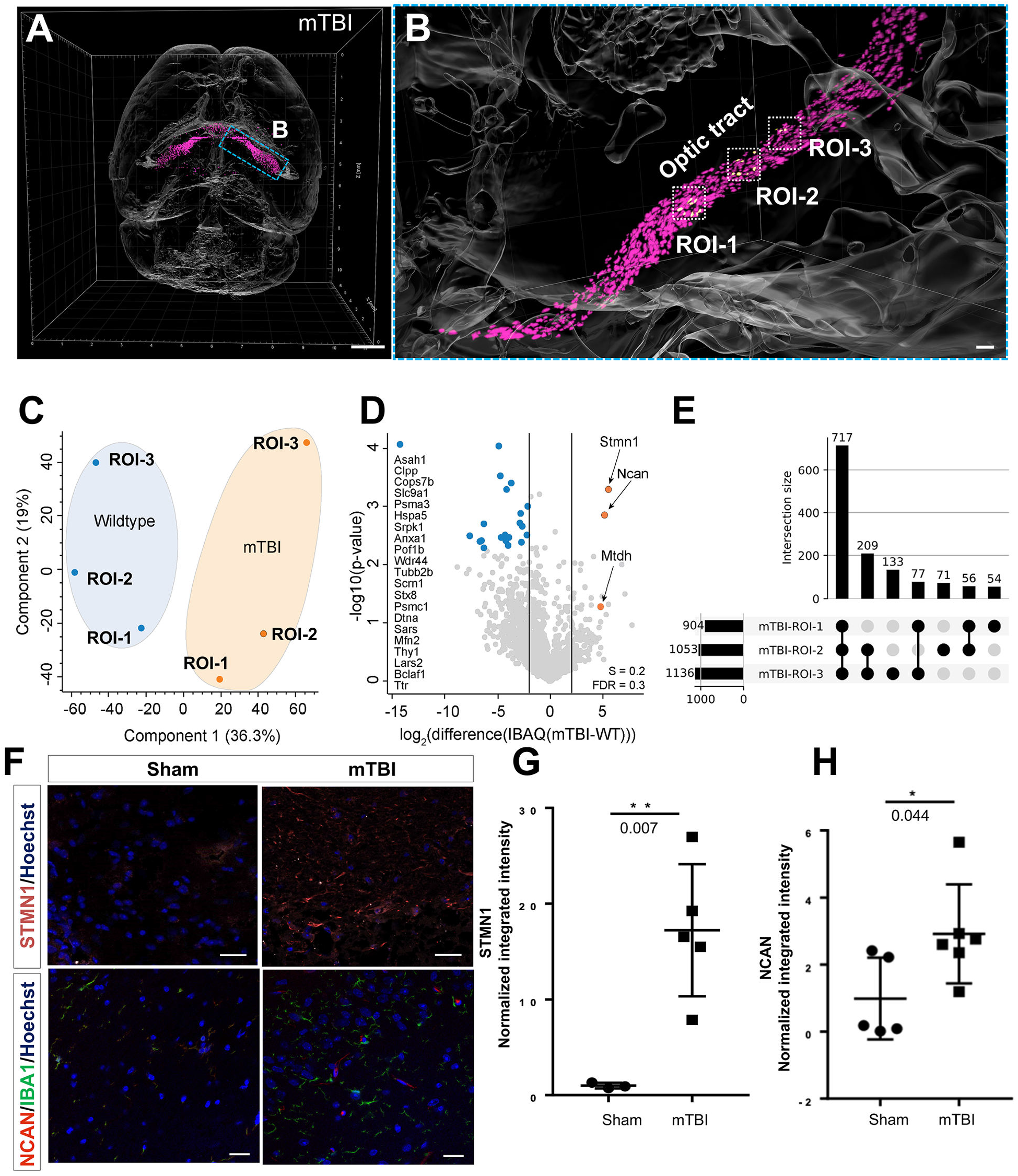
DISCO-MS reveals effects of mTBI in discrete regions of whole brain. **(A)** 3D-reconstruction of an exemplary CX3CR1^GFP/+^ mouse brain after mTBI. Segmented microglia shown in magenta. Scale bar, 500 μm **(B)** High-magnification view of region shown in (A), highlighting substantial increase in activated microglia along optic tract region. 3 neighboring ROIs along optic tract were identified, laser captured and subjected to proteomic analyses. Scale bar, 100 μm **(C)** PCA plot showing the distribution of individual ROIs from mTBI vs. ROIs from control (sham surgery) with the same spatial location. **(D)** volcano plot showing the significant enrichment of proteins. **(E)** The number of shared and unique set of proteins in ROIs in mTBI. **(F)** Histological validation of top 2 proteins in optic tract: 1) stathmin (STMN1) shown in red and nuclear marker Hoechst dye in blue; 2) neurocan (NCAN) shown in magenta along with microglia marker (IBA1) in green and Hoechst dye in blue. Scale bars, 20 μm **(G, H)** Intensity quantification of STMN1 immunostaining signal (*P* = 0.007, n=3 animals per group from total 9 sections) and NCAN immunostaining signal (*P* = 0.044, n=3, animals from total 11 sections) in mTBI vs. sham controls, respectively (unpaired two-sided Student’s t-test, data presented as ± SD).

**Fig. 4:**
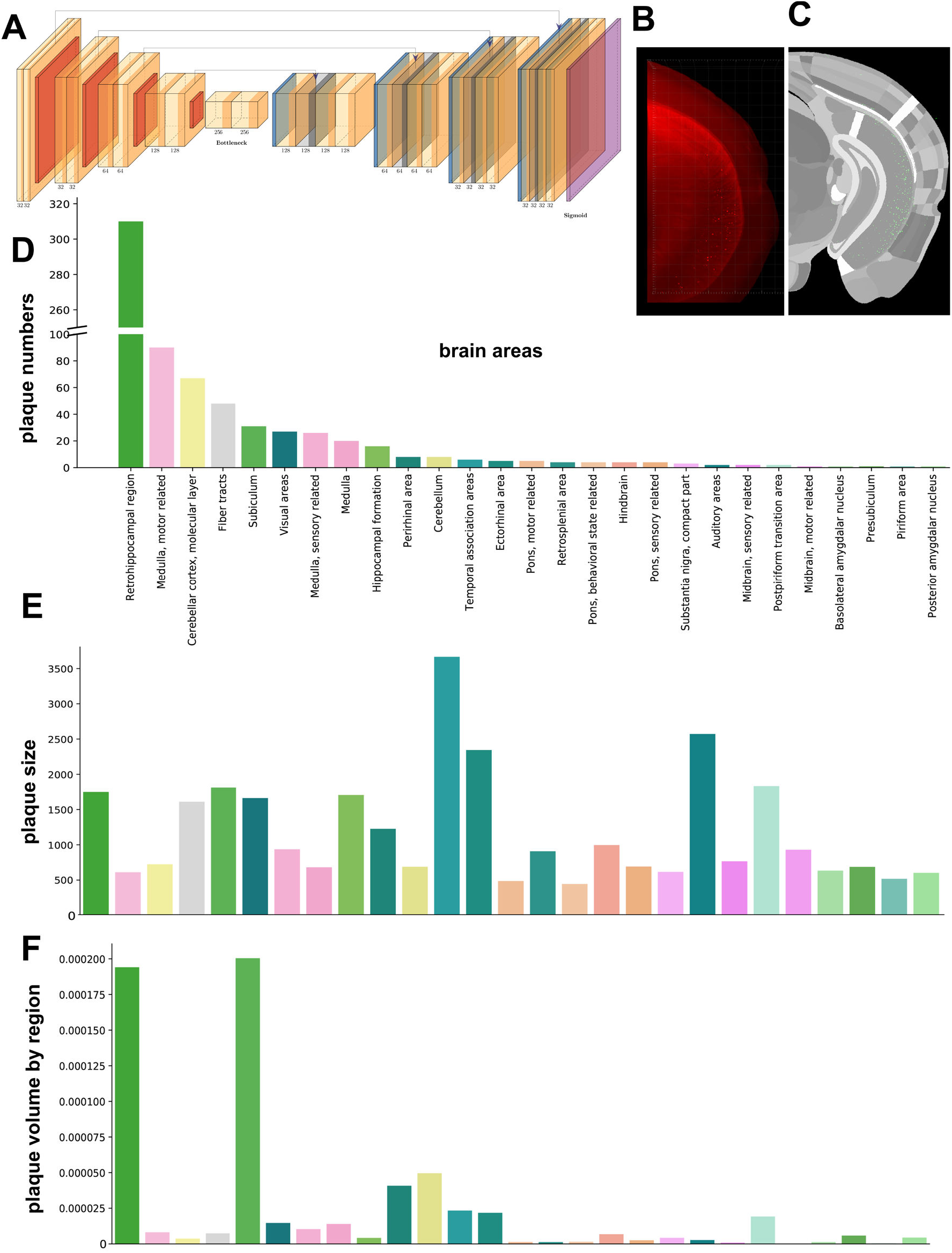
Deep learning analysis of plaques in whole 5×FAD mouse brains. **(A)** 3D U-Net architecture including layer information and feature sizes. **(B)** Maximum intensity projection of the hippocampal region of the brain **(C)** segmented plaques overlaid on the hierarchically and randomly color-coded atlas to reveal all annotated regions available. After deep learning-based quantification, the data was registered to Allen brain atlas to obtain region-wise quantification. **(D)** the number of the plaques in the major brain areas. **(E)** the average plaque size per brain region. **(F)** the density of plaques: the ratio of plaque volume per brain region volume.

### Scalable and robust pathology identification using deep learning

One of the early hallmarks of Alzheimer’s disease (AD) pathology is the accumulation of amyloid-beta (Aβ) plaques in the brain parenchyma (Meyer-Luehmann et al., 2008). We imagined that unbiased detection of all Aβ plaques, followed by their equally unbiased proteome analysis using DISCO-MS would provide valuable insights to the initial stages of AD. To explore this hypothesis, we used the 5×FAD mouse model of AD and aimed for the identification of Congo red-labelled Aβ plaques in young mouse brains.

As the locations of these initial plaques are unknown, we developed a deep learning approach to identify all Aβ plaques rapidly and reliably in whole mouse brain scans. In short, our network architecture is based on U-Net, a well-established approach for biomedical image analysis (Ronneberger et al., 2015). As the loss function, we used an equally weighted combination of Dice and binary cross entropy, and the Ranger optimizer, which combines Rectified Adam, gradient centralization and LookAhead (**Fig. 4A**). We applied suitable data augmentation protocols for training and implemented test time augmentation. To assess our segmentation quality, we calculated a wide range of voxel-wise and Aβ plaque segmentation metrics. Our deep learning architecture for automated Aβ plaque detection showed high performance in volumetric accuracy (0.99±0.00), volumetric (0.71±0.06) and surface (0.94±0.03), as well as overall Dice scores (0.89±0.09) per Aβ plaque. After segmenting all plaques in the entire brain using deep learning, we registered our data to the Allen brain atlas to obtain region-wise quantifications for more than a thousand brain subregions (**Fig. 4B,C**) (Wang et al., 2020; Todorov et al., 2020). We then grouped them into the major brain regions defined by the Allen mouse brain ontology for the simplicity.

Our deep learning model identified Aβ plaques already in six weeks old mouse brains, much earlier than any previous study suggested (Boza-Serrano et al., 2018). Some of the main brain regions with initial plaques were retro hippocampal region (310 plaques), medulla (motor area, 90 plaques), molecular layer of cerebellar cortex (67 plaques), fiber tracts (48 plaques), subiculum areas (31 plaques), visual area (27 plaques) and hippocampal formation (16 plaques) (**Fig. 4D**). We also analyzed the volume of detected Aβ plaques in these brain areas and observed the largest plaques in the temporal association, ectorhinal and auditory areas with an average volume between 2000-3500 μm^3^ (**Fig. 4E**). Moreover, brain regions with the highest density of Aβ plaques were retro hippocampal and the subiculum areas (**Fig. 4F**).

After deep learning-based identification of Aβ plaques in the 5xFAD mouse model, we isolated four ROIs (volume: ~0.0005 mm^3^) from the hippocampal region (vs. corresponding brain regions from the control mice) and subjected them to MS-based proteomics (**Fig. 5A–D**). We also confirmed our finding of Aβ plaques in six-week-old mice in the same brain regions by immunohistochemistry with anti-amyloid beta monoclonal antibodies (**Fig. S5A**). We compared ~2000 proteins across replicates and PCA plot separated the ROIs with Aβ plaques from the control brain regions (**Fig. 5E**). Differential expression analysis revealed that many well-characterized AD-associated proteins were enriched in 5xFAD ROIs including the amyloid-beta precursor protein (Meyer-Luehmann et al., 2008; Lichtenthaler, 2006) (32-fold increase) and the thimet oligopeptidase 1 (8-fold increase) (**Fig. 5F**). Apart from known and well-established AD-related proteins (Meckelein et al., 1996; Pollio et al., 2008), we also detected less characterized proteins in early-stage Aβ plaques such as a member of the calcium-binding protein family S100a11. Additional experiments highlighted the absence of Aβ plaques at five weeks of age, whereas plaques were evident at seven weeks (**Fig. S5B**).

**Fig. 5:**
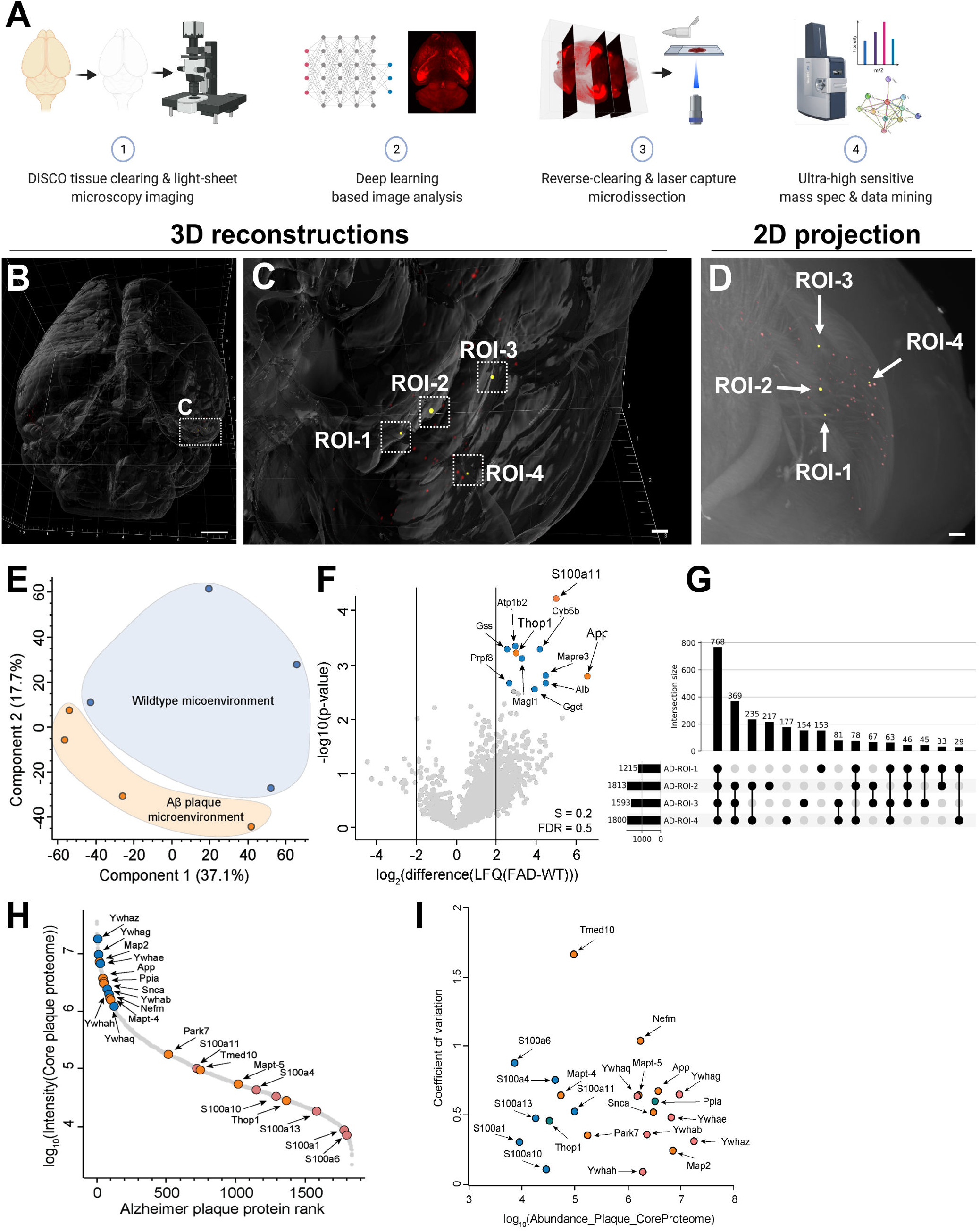
DISCO-MS unravels the single plaque proteome in AD mouse model. **(A)** Major steps of DISCO-MS (See also Movie S1). **(B)** 3D visualization of all Aβ plaques (in red) in 5×FAD mouse brains at 6 weeks age (n=4 experimental replicates). Scale bar, 500 μm **(C)** Enlarged view of marked region in **(B)**, 4 different ROIs from hippocampus - each containing single plaque-selected and isolated for mass spectrometric measurements. Scale bar, 100 μm **(D)** 2D projection of selected ROIs. Scale bar, 100 μm **(E)** PCA plot of ROIs’ proteome from 5xFAD vs. the same regions from wildtype controls. **(F)** Volcano plot showing the significantly enriched proteins. **(G)** The number of shared and unique set of proteins in 5×FAD ROIs. **(H)** Rank order of core protein signals in a single plaque microenvironment. **(I)** Log_10_ abundance distribution of selected proteins and protein families as a function of their coefficient of variation (CV) across the core Aβ plaque proteomes. Dynamic range coverage is up to four orders of magnitude. CVs indicate variability in the shared plaque core proteome, among proteins known to play a role in Alzheimer’s disease.

Moreover, we asked how similar the proteome in our ROIs with early Aβ plaques are. Plaques with more than 1200 protein identifications each shared a total of 768 proteins, defining the core proteome of early-stage Aβ plaque formation (**Fig. 5G)**. An abundance rank plot of the shared early-stage Aβ plaques core proteome revealed several members of the Ywhaz (14-3-3) and the S100a protein family. We also found many other proteins known to be involved in AD such as Mapt, Snca and Park7. Illustrating the specificity of MS-based proteomics, we identified two isoforms of Mapt, namely Mapt-4 (Tau C) and Mapt-5 (Tau D) in the early-stage Aβ plaque ROIs (**Fig. 5H**). Our proteomics data also suggest early-stage Aβ plaque variability (**Fig. 5**I) with regards to well-characterized AD proteins (Mapt, Tmed10, Park7, Snca, App), proteins of the S100a family (a6, a4, a13, a1, a10, a11), peptidases (Thop1, Ppia), proteins of the Ywha family (14-3-3, q, g, e, b, h, z) and other structure-determining proteins including Nefm and Map2. Finally, we confirmed the presence of S100a11 and Thop1 in early-stage plaques of 5xFAD brain slices by immunofluorescence (**Fig. 6**). Thus, DISCO-MS recovered many known markers of Aβ plaques in AD and revealed potentially novel ones. Our data also suggest that S100a family members could be another driving factor in early-stage Aβ plaque development (Cristóvão and Gomes, 2019). Taken together, DISCO-MS allowed us to pinpoint early Aβ plaques in whole brains and analyze the proteomic makeup of isolated regions of interest including identified Aβ aggregates (**Movie S1**).

**Fig. 6:**
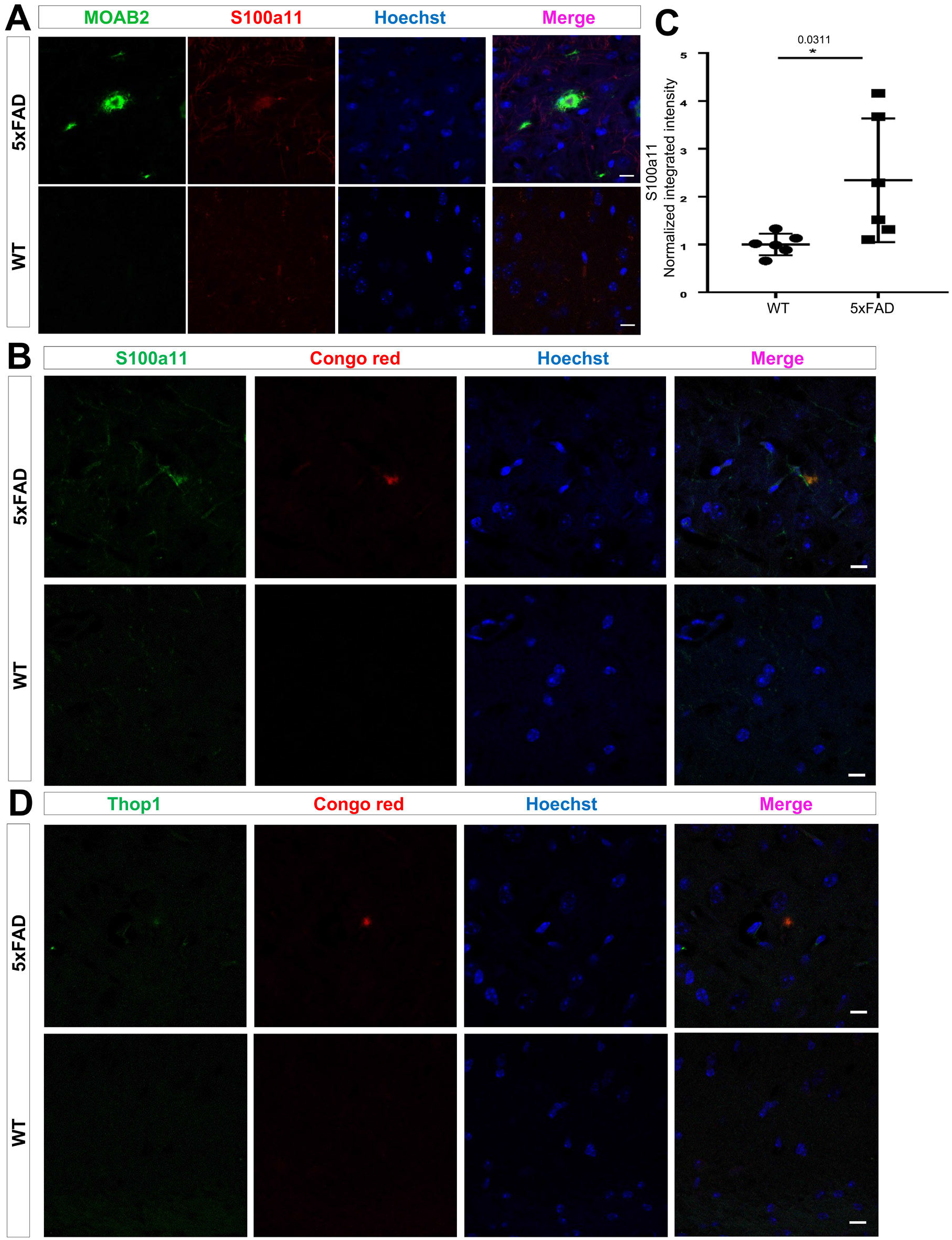
Histological validation of DISCO-MS hits. **(A)** Histological validation of DISCO-MS hit S100a11 (red) in hippocampal region using antibody immunostaining in 6 weeks old mice. The plaques were co-labeled using the MOAB2 antibody (in green). **(B)** We confirmed the S100a11 signal also around Congo red-labeled plaques in 6 weeks old mice. **(C)** Intensity quantification of S100a11 (N=3 animal per group from total 12 sections; unpaired two-sided Student’s t-test; p = 0.0311; data are presented as average ± SD). **(D)** Histological validation of DISCO-MS hit Thop1 (in green) in hippocampal region of 5xFAD animals along with Congo red-labeled (in red) plaques in 6-weeks old mice. Scale bars, 10 μm

## DISCUSSION

Deciphering tissue heterogeneity is essential to better understand developmental and pathological processes. Enormous progress has been made in single-cell transcriptomic technologies throughout the last years (Ji et al., 2020; Stuart and Satija, 2019; VanHorn and Morris, 2021). Furthermore, novel ultra-high sensitivity MS-based approaches have emerged, enabling the analysis of proteomes down to the level of several labeled and pooled or even true single cells (Brunner et al., 2020; Budnik et al., 2018; Cheung et al., 2021; Cong et al., 2021; Mund et al., 2021). A major leap for both technologies would integrate proteomics and transcriptomics data with their spatial information (Rodriques et al., 2019; Wang et al., 2018). Towards this goal, new technologies emerged for the molecular and spatial characterization on tissue slices (Alon et al., 2021; van den Brink et al., 2020; Chen et al., 2020; Liu et al., 2020; Mahdessian et al., 2021; Rodriques et al., 2019; Ståhl et al., 2016; Takei et al., 2021). In parallel, tissue clearing methods enabled imaging cellular even sub-cellular level details of protein/ molecule of interests in intact transparent organs and organisms (Yang et al., 2014; Ueda et al., 2020; Cai et al., 2019; Blutke et al., 2020; Richardson and Lichtman, 2015).

As many pathologies start in yet unknown regions of organs, a method first to determine the whereabouts of initial tissue abnormalities in whole organs, then to reveal their complete molecular profiles would be essential in understanding and handling diseases at their early stages. We addressed this challenge by developing and applying DISCO-MS for bottom-up proteome analysis of tiny tissue regions identified by panoptic imaging of intact organs and in silico 3D reconstructions using AI. Using DISCO-MS, we identified initial brain regions that form Aβ plaques in the brains of young AD mouse model. Expectedly, most of the initial plaques appeared in the Retrohippocampal regions, including the Entorhirnal area, and surprisingly some in hindbrain regions, including Pons and Medulla (Whitesell et al., 2019). As our work also provides proteome data, it can facilitate the discovery of new diagnostic and therapeutic approaches tailored for the early stages of AD. Our technology performs equally well on both rodent and human tissues and yields qualitative and quantitative proteomics data nearly indistinguishable from uncleared samples, even for the harshest organic solvent-based tissue clearing approaches. Although tissue clearing presumably removes the lipidic cast of membrane proteins, we observed that the plasma membrane protein gene ontology class was hardly affected, suggesting that DISCO-MS could be used to identify novel surface markers for drug targeting. Beyond neuroscience, this technology can transform spatial molecular investigations in many other biomedical research areas, among others revealing molecular profiles of cancer micrometastases in whole mouse bodies (Pan et al., 2019) and human biopsies, as well as invasion mechanisms of pathogens and viruses, including SARS-CoV-2 in both in animal models and human organs (Sun et al., 2020; Trypsteen et al., 2020).

Notably, the DISCO-MS technology presented here is versatile across labeling and solvent cleared methods for whole organs. It is not only applicable to reporter mouse lines but can also be utilized where reporter lines are currently absent. In those cases, deep tissue labeling with antibodies and/or dyes can be performed against the antigen of interest, imaged as a whole organ, and subjected to MS-based proteome profiling. DISCO-MS should be of great interest to researchers in already possession of archived solvent cleared organs and imaging data, where molecular data is missing. Now, this method will further enable understanding the molecular basis of a pathological milieu in these organs. In conclusion, we present a spatial unbiased proteome profiling technology comprising complete 3D imaging data of whole organs, enabling unbiased identification of interesting tissue regions for subsequent molecular characterization.

## ACKNOWLEDGMENTS

A.-D.B. acknowledges support from the International Max Planck Research School for Life Sciences-IMPRS-LS. We thank our colleagues in the Department of Proteomics and Signal Transduction, Max Planck Institute of Biochemistry, for discussions and help. In particular, we thank I. Paron, A. Piras and C. Deiml for technical support and J.B. Müller for column production, and I. Bechmann and H. Steinke for human tissues. S.F.L acknowledges the receipt of funding from Deutsche Forschungsgemeinschaft (DFG, German Research Foundation) under Germany’s Excellence Strategy within the framework of the Munich Cluster for Systems Neurology (EXC 2145 SyNergy– ID 390857198) and M.M. & S.F.L. by the Federal Ministry of Education and Research through grant CLINSPECT-M (FKZ161L0214C).

## AUTHOR CONTRIBUTIONS

H.S.B., A.-D.B., M.M. and A.E. conceptualized and designed the study. H.S.B., A.-D.B., Z.R, H.M, M.T., M.M, Z.I.K performed experiments. H.S.B, Z.R, H.M performed mice experiments, solvent-based organ clearing, light sheet imaging procedures and stitching of data. H.S.B. developed the LCM-based isolation procedure for target tissues from solvent-cleared organs. A.-D.B. developed the sample preparation workflow for proteomics analysis. A.-D.B. and M.T. performed mass spectrometry-based proteomics analysis. A.-D.B. performed proteomics data analysis. R.A., J.C.P., F.K., M.A., B.H.M. and F.J.T. developed deep learning models. M.A., F.J.T. performed data analysis. M.I.T. performed atlas registration of 5xFAD brains. H.S.B. and D.K. performed ClearMAP analyses. S.M. and S.F.L. helped with the prototyping experiment. H.S.B., A.-D.B., M.T., analyzed the data. H.S.B., A.-D.B., M.M. and A.E. wrote the manuscript.

## COMPETING FINANCIAL INTERESTS

F.J.T. reports receiving consulting fees from Roche Diagnostics GmbH and Cellarity Inc., and ownership interest in Cellarity, Inc. and Dermagnostix. All other authors have no competing interests.

## DATA AVAILABILITY

All mass spectrometry raw data, libraries and outputs from each particular search engine analyzed in this study have been deposited to the ProteomeXchange Consortium via the PRIDE partner repository and made available to the reviewers.

## METHODS

### Animals

We used the following animals in the study: mixed gender CX3CR1-eGFP, Thy-1-GFPM, 5xFAD and C57Bl6/J from Jackson Laboratory. The animals were housed under a 12/12 hours light/dark cycle. The animal experiments were conducted according to institutional guidelines: Klinikum der Universität München / Ludwig Maximilian University of Munich and after approval of the Ethical Review Board of the Government of Upper Bavaria (Regierung von Oberbayern, Munich, Germany) and the Animal Experiments Council under the Danish Ministry of Environment and Food (2015-15-0201-00535) and following the European directive 2010/63/EU for animal research. All data are reported according to the ARRIVE criteria. Sample sizes were chosen based on prior experience with similar models.

### Human samples

Intact human brains were taken from human body donors with no known neuropathological diseases. The donors gave their informed and written consent to explore their cadavers for research and educational purposes, when still alive and well. The signed consents are kept at the Anatomy Institute, University of Leipzig, Germany. Institutional approval was obtained in accordance to the Saxonian Death and Funeral Act of 1994. The signed body donor consents are available on request.

### Repeated closed head mild traumatic brain Injury (mTBI)

Before mTBI, tin foil was taped and tightened to the U-shaped stage made of clear plastic container (38 x 27 x 27 cm^3^) containing a sponge collection (38 × 25 × 15 cm^3^). Then, mice were pre-treated with buprenorphine (1:15 saline, 50 ul/20 mg, ip) and anesthetized with 4% isoflurane using 1.0 L min^−1^ air until non-responsive to a paw or tail pinch. To ensure the head acceleration and rotation following the head impact, the mice were placed under impact tip on the tin foil, which contains holes according to the shape of a mouse and can support the body weight of the mice. The mice were kept under light anesthesia with continued 2% isoflurane. mTBI was produced using a stereotaxic impactor device with a 5-mm round tip coated with 1mm thick rubber, which can preserve an intact skull after impact. Impact tip was placed and covered the scalp’s area from just behind the eyes to the midline of the ears, the center of the tip at approximately midway along the sagittal suture. The injury was produced without skin incision (velocity of 5 m/sec, depth of 0.5 mm, and dwell time of 0.1 sec). The mouse would be removed quickly from the collection sponge and transferred to the recovery box maintained at 32 °C. In total, mice received four hits with a 48-hour interval in seven days. Sham mice receive identical handling and exposure to the same time length of anesthesia as the mTBI mice but received no impact.

### Perfusion and tissue preparation

Mice were anesthetized using a combination of midazolam, medetomidine and fentanyl (MMF) (1mL/100g of body mass for mice; i.p.). As soon as the animals did not show any pedal reflex, they were intracardially perfused with cold heparinized 0.1 M PBS (10 U/mL of Heparin, Ratiopharm; 100-125 mmHg pressure using a Leica Perfusion One system) for 5-10 minutes at room temperature until the blood was washed out, followed by ice-cold 4% paraformaldehyde (PFA) in 0.1 M PBS (pH 7.4) (Sigma) for 10 minutes. Then, the brains were extracted and post-fixed in 4% PFA for 1 day at 4 °C and later washed with 0.1 M PBS for 10 minutes 3 times at room temperature. The whole brain clearing or nanoboosting procedure was started immediately. For the collection of fresh frozen samples, animals were sacrificed by cervical dislocation and brains were quickly snapped frozen in liquid nitrogen and stored in −80 °C until further processing.

### Congo red labeling of whole brains of 5×FAD animals

Whole brains were dehydrated with gradual addition of methanol in PBS (50% x1, 80% x1, 100% x2, each for 1 hr). Overnight bleaching with 5% hydrogen peroxide in methanol was done at 4 °C. Brains were then gradually rehydrated in 100%, 80%, 50% methanol in PBS (1 hr for each step, followed by 2 additional washes in PBS). Detergent washing was then performed in PBS with 0.2% Triton X-100 for 2 hr, brains were incubated overnight at 37°C in PBS with 0.2% Triton X-100 and 0.3 M glycine, followed by blocking in PBS with 0.2% Triton X-100 and 6% goat serum for 7 days. Following blocking, the tissue was washed for 1 hr twice in PBS with 0.2% Tween-20 and 10 μg/ mL heparin (PTwH). Next, brains were incubated with 10 μM Congo Red (Sigma, C6277) at 37°C in PTwH for 5 days. After that brains were washed in PTwH for 2 days with periodic solution changes and gradually dehydrated using 3DISCO clearing as described next.

### Clearing of brains using DISCO methods

We followed the 3DISCO and uDISCO passive clearing protocol as described previously. In brief, dissected brains were placed in 5 ml tubes (Eppendorf, 0030 119.401) and covered with 4.5 mL of clearing solution. All incubation steps were performed in a fume hood with gentle shaking or rotation, with the samples covered with aluminum foil to keep them in dark. To clear the samples using 3DISCO, gradient of tetrahydrofuran (THF) in distilled water (vol/vol%), 2 hours for each step, was used as 50%, 70%, 90%, 100% and overnight 100 % THF; after dehydration, the samples were incubated for 45 min in dichloromethane (DCM, Sigma, 270997), and finally in BABB (benzyl alcohol + benzyl benzoate 1:2, Sigma, 24122 and W213802) until transparency. Next for uDISCO a gradient of tert-butanol (Sigma, 360538) in distilled water (vol/vol %) was used as 50%, 70%, 90%, 100% twice at 32°C for 12 hours each step, followed by immersion in DCM for 45 minutes at room temperature and finally incubated with the refractive index matching solution BABB-D15 containing 15 parts BABB, 1 part diphenyl ether (DPE) (Alfa Aesar, A15791) and 0.4% Vol vitamin E (DL-alpha-tocopherol, Alfa Aesar, A17039), for at least 6 hours at room temperature until achieving transparency.

### vDISCO whole-brain passive immunostaining, clearing and imaging

Passive vDISCO was performed on dissected organs as performed by Cai R et al. First, the post-fixed brains were pretreated with permeabilization solution containing 1.5% goat serum, 0.5% Triton X-100, 0.5 mM of Methyl-beta-cyclodextrin, 0.2% trans-1-Acetyl-4-hydroxy-L-proline and 0.05% Sodium Azide 0.1 M for 2 days at 37°C with gentle shaking. Subsequently, the brains were incubated in 4.5 mL of this same permeabilization solution plus the nanobooster Atto647N conjugated anti-GFP (1:600, which is ~5-8 μg of nanobooster in 4.5 ml) for CX3CR1-eGFP and Thy-1-GFPM brains for 12-14 days at 37°C with gentle shaking, then brains were washed for 2 hours 3 times and once overnight with the washing solution (1.5% goat serum, 0.5% Triton X-100, 0.05% of sodium azide in 0.1 M PBS) at room temperature and in the end washed for 2 hours 4 times with 0.1 M PBS at room temperature. The immunostained brains were cleared with 3DISCO clearing first they were put in the Eppendorf 5 ml tubes and then incubated at room temperature with gentle shaking in 4.5 mL of the following gradient of THF in distilled water (vol/vol%), 2 hours for each step: 50%, 70%, 90%, 100% THF and overnight 100 % THF; after dehydration, the samples were incubated for 45 min in DCM, and finally in BABB until transparency. During all the clearing steps, the tubes were wrapped with aluminum foil to keep them in dark.

### SHANEL sample preparation and clearing

Archived human brain samples were obtained in PFA which were stored for a long period of time (>5 years) at 4 °C and subjected to our previously published SHANEL clearing protocol with some modifications (Zhao et al., 2020). Briefly, samples were dehydrated with EtOH/dH_2_O series at RT: 50%, 70%, 100% for 1 h for each step. Then incubated with 10 mL DCM/MetOH (2:1 v/v) (freshly prepared) for 6 h at RT followed by rehydration with EtOH/dH_2_O series at RT: 100%, 70%, 50%, dH_2_O for 1 h each step then incubated with 0.5M acetic acid (30 mL/L) at RT for 2 h, then wash with dH_2_O twice for 15 min and then incubated with 4M guanidine hydrochloride (382.12 g/L), 0.05M sodium acetate (4.1g/L), 2% v/v Triton X100 in dH_2_O, (measure pH: 6.0) at RT for 2 h, then wash with dH_2_O twice for 15 min each and wash with PBS twice for 15 min each. Afterwards samples were incubated with 10% CHAPS, 25% N-Methyldiethanolamine in dH_2_O at 37 °C for 4 hours and then washed with dH O twice for 15 min each. Since we did not perform any deep antibody labeling in these samples, we started clearing these samples without prior blocking or antibody labeling steps. Clearing was done with THF in water with dilutions (vol/vol %) of 50%, 70%, 90%, 100%, 1hr each, 100% overnight, DCM 45 min and incubated in BABB until the samples were transparent.

### Behavioral assessment

#### Barnes Maze

Briefly, a maze consisting of a surface bright circular platform with an escape black box can be recessed and located at the bottom of one of the 20 holes. Visual shapes were placed on 3 walls of the room as cues. For all trials, mice were placed in a cylinder black start chamber in the center of the maze for 10 second. After the chamber lifted and the test started, mice were given 3 minutes to locate and enter the target box during the spatial acquisition time. For a period of 4 days, 4 trials were given per day with an inter-trial of 15 minutes. The trial ended when the mouse entered the escape box or after 3 minutes had elapsed. Mice were allowed to remain in the escape box for 1 minute. A system (Ethovision XT) was used to continually track and record the movement of the mice. Escape latency was measured as the time taken for the mouse to enter the box.

#### Assessment of motor function

The mice were given 3 trials training per day for 3 days to walk along a 1 cm diameter and 100 cm long wood beam with a goal box on the end of the beam before mild TBI. The height of the beam was placed 1 m above ground. The latency that it takes walking to cross the beam after 8 weeks mild TBI were recorded. Mice that were unable to cross the beam would be removed in the training.

### Immunofluorescence and Confocal microscopy

Briefly, mice were sacrificed after 8 weeks of injury or at six weeks of age following transcardial perfusion with PBS and with 4% cold PFA. Brains were post-fixed in 4% PFA at 4 °C overnight. Either frozen sections or cleared-rehydrated frozen sections were treated with 0.2% Triton X-100 in PBS for 15 min, blocked for 1 hour at room temperature with 10% serum in PBST. Then incubation with primary antibodies Iba1 (1:1000, Wako, 019-19741), Stathmin 1 (1:300, Novus, NBP1-76798), Neurocan (1:300, abcam, ab31979), S100a11 (1:300, R&D, MAB5167), Thop1 (1:300, Novus, NB400-146), MOAB2 (1:1000, Novus, NBP2-13075), at 4°C for overnight and Alexa conjugated secondary antibodies (1:1000, Goat anti-rabbit IgG Alexa fluor 647, Invitrogen, A21245; Goat anti-Mouse IgG Alexa Fluor 488, A11029; Goat anti-Mouse IgG Alexa Fluor 594, A11032; Goat anti-Rat IgG Alexa Fluor 594, A11007; Goat anti-Rat IgG Alexa Fluor 488, A11006;) were incubated for 1 hour at room temperature. Slices were mounted after being stained with Hoechst 33342 (Invitrogen). Images were acquired with 10x, 40x, and 63x objective of confocal microscope (ZEISS LSM880).

### Light-sheet microscopy and image processing

Single plane illuminated (light-sheet) image stacks were acquired using an Ultramicroscope II (LaVision BioTec), featuring an axial resolution of 4 μm with following filter sets: ex 470/40 nm, em 535/50 nm; ex 545/25 nm, em 605/70 nm; ex 640/40 nm, em 690/50 nm. Whole brains were imaged individually using high magnification objectives: 4x objective (Olympus XLFLUOR 4x corrected/0.28 NA [WD = 10 mm]), LaVision BioTec MI PLAN 12x objective (0.53 NA [WD 10 = mm]) coupled to an Olympus revolving zoom body unit (U-TVCAC) kept at 1x. High magnification tile scans were acquired using 20% overlap and the light-sheet width was reduced to obtain maximum illumination in the field. Processing, data analysis, 3D rendering and video generation for the rest of the data were done on an HP workstation Z840, with 8 core Xeon processor, 196 GB RAM, and Nvidia Quadro k5000 graphics card and HP workstation Z840 dual Xeon 256 GB DDR4 RAM, nVidia Quadro M5000 8GB graphic card. We used Imaris (Bitplane), Fiji (ImageJ2) and Vision 4D (Arivis) for 3D and 2D image visualization. Tile scans were stitched by Fiji’s stitching plugin49.

### ClearMap quantification

To quantify microglia distribution in whole brains of mTBI and sham animals, we used ClearMap. As the script was originally developed for quantification of the cFos+ cells, to comply with the offered method, we did the following pre-processing steps on our microglia data using Fiji before ClearMap:

1. Background equalization to homogenize intensity distribution and appearance of the microglia cells over different regions of the brain, using pseudo-flat-field correction function from Bio-Voxxel toolbox.
2. Convoluted background removal, to remove all particles bigger than relevant cells. This was done with the median option in the Bio-Voxxel toolbox.
3. Two-dimensional median filter to remove remaining noise after background removal. The filter radius was chosen to ensure the removal of all particles smaller than microglia cells.
4. Unshapen mask to amplify the high-frequency components of a signal and increase overall accuracy of the cell detection algorithm of ClearMap.

After pre-processing, ClearMap was applied by following the original publication and considering the threshold levels that we obtained from the pre-processing steps. As soon as the quantification was completed, the data was exported as an Excel file for further analysis.

### Deep learning analyses

The segmentation of the stained Aβ plaques represents a key step towards a reliable quantification thereof. We develop a customized, three-dimensional deep learning approach to optimize segmentation of Aβ plaques in the whole brains of 5×FAD animals. Our network architecture is inspired by the well-established U-Net architecture. The used loss function is an equally weighted combination of Dice and binary cross entropy loss. We use the Ranger optimizer, which combines Rectified Adam, gradient centralization and LookAhead. Our dataset consists of 98 image volumes (300×300×300 voxel) from one Alzheimer brain; where 34 volumes include Aβ plaques and 64 volumes do not contain any plaques. An ensemble of experts including the scientist who imaged the brains labeled all images. We randomly sample our training set of 85 volumes (21 with AD plaques, 64 without AD plaques), our validation set of seven volumes and our separate test set of six volumes. During training and testing we applied suitable data augmentation protocols. In order to assess the quality of our segmentation we calculate a wide range of voxel-wise and Aβ plaque wise segmentation metrics. Based on the reliable segmentation of individual Aβ plaques we continue towards a statistical evaluation of the number and size of Aβ plaques per brain region. First, we register all of our brains to the Allen brain Atlas, enabling a single voxel assignment to brain structures. For whole brain segmentation we extract the single connected components, which represent our individual segmented Aβ plaques and calculate their total size in voxels as a biomarker. Using this registration and biomarker, we calculate per brain region statistics for the presence and size of Aβ plaques across the whole brain.

### Optimization of cleared tissue for cryopreservation and sectioning

After acquiring the whole brain images from CX3CR1-eGFP, 5xFAD, C57BL/6J mice the brains were further optimized for cryopreservation and sectioning. The course of tissue clearing and imaging in BABB makes the tissue brittle and hard for further processing, Thus, to solve this hurdle we re-hydrated the samples with respective clearing solutions to be able to process samples for cryosectioning. Thereafter, samples were washed with PBS twice for 15 min each and cryopreserved overnight with 30% sucrose solution in 4 °C. In order to avoid any ice crystals formation, samples were further embedded in Optimal cutting temperature compound (OCT compound) under the chilled isopentane container placed on dry ice. Samples were stored in −80 °C until cryosectioning.

### Laser-capture microdissection

For the microdissection of cells and plaques, we used the PALM MicroBeam system (Zeiss). PALM MicroBeam uses a focused laser beam to cut out and isolate the selected specimen without contact. The laser catapult isolates the region of interest fast and uncontaminated in the adhesive cap mounted in the RoboMover. Briefly, after cryosectioning, sections were mounted on the polyethylene naphthalate (PEN, Zeiss) slides and were either stored at −80 °C in 50 ml falcon tubes filled with molecular sieves (Sigma-Aldrich) or processed further for serial dehydration with ethanol and air dried for 15 min under the hood. Cells in the optic tract from mTBI/Sham brains and amyloid-beta plaques from the 5xFAD and from respective WT brain regions were microdissected by laser pressure catapulting (LPC) UV laser capture microdissection system (Palm Zeiss Microlaser Technologies, Munich, Germany) consisting of an inverted microscope with a motorized stage, an ultraviolet (UV) laser and an X-Cite 120 fluorescence illuminator (EXFO). The microdissection process was visualized with an AxioCam ICc camera coupled to a computer and was controlled by Palm RoboSoftware (Zeiss, Germany). Approximately an area of 200×200 μm (corresponding to 40-60 cells) were cut by laser using 20x objective (LD Plan-Neofluar 20x/0.4 corr M27) and catapulted against gravity into the adhesive cap. Tissues were quickly lysed in 20 μl of lysis buffer, spinned down and kept in dry ice or stored in −80 °C. To avoid any uncertainties in capturing ROI, each time after catapulting as well as after lysing and spin down, adhesive cap was properly examined under the camera.

### Optimization of DISCO cleared sample preparation for mass spectrometry analysis

Several conditions and combinations of solubilizing agents for the isolation of proteins from tissue cleared mouse brain, heart, and lung samples were initially evaluated for protein extraction efficiency, peptide recovery, and qualitative and quantitative reproducibility keeping fresh or PFA-fixed as reference. Our goal was to establish a workflow that recovers proteomes that are as similar as possible to non-cleared tissue and is universal for all tissue clearing techniques.

Cleared organs or cryosections were removed from the refractive index matching solution BABB and washed five times with 1x PBS solution. The organ was then flash-frozen and pulverized in a Covaris CP02. Afterwards, the samples were resuspended in different protein solubilizing solutions (6% Sodiumdodecylsulfate, 500 mM TrisHCl, pH 8.5 (SDS buffer); 2% Sodiumdeoxycholate, 100 mM TrisHCl pH 8.5, 10 mM Tris-(2-carboxyethyl)-phosphin (TCEP), 40 mM Chloroacetamide (SDC buffer); 50% Trifluoroethanole, 100 mM TrisHCl, pH 8.5 (TFE buffer), followed by protein extraction at 95 °C, 1.000rpm for 45 min. Then the samples were subjected to sonication (Branson) at maximum frequency for 30 cycles at 50% output, followed by another heating step at 95 °C, 1.000 rpm for 45 min. From here on, processing steps diverged for each protocol.

Proteins solubilized in the SDS buffer were precipitated with ice-cold Acetone at 80% v/v ratio overnight at −80 °C, followed by centrifugation at max. g for 15 min at 4 °C. The supernatant was removed, the pellet was washed with 5 ml ice-cold 80% v/v Acetone/ddH_2_O, followed by 30 min precipitation on dry ice. The acetone wash steps were repeated two times for a total of three washes. Proteins solubilized in the TFE buffer, were subjected to solvent evaporation in a speedvac at 45 °C until dryness before further processing.

In case of SDS-SDC or TFE-SDC protocol, in which SDS or TFE protein extraction was coupled to an SDC-based protein digestion, SDS- or TFE-solubilized proteins were resuspended in 1ml of SDC buffer and heated to 95 °C at 1.000 rpm for 10 min to denature proteins, reduce cysteine bridges and alkylate free cysteine residues. Afterwards, samples were sonicated for 15 cycles each 30sec at max power in a Bioruptor, followed by another heating step for 10 min at 95 °C, 1.000 rpm in a Thermoshaker.

SDC-only, SDS-SDC, TFE-SDC solubilized protein solutions were cooled down to room temperature, diluted 1:1 with 100 mM TrisHCl, pH 8.5, followed by protein concentration estimation by Nanodrop. Extracted and solubilized proteins were digested overnight at 37 °C and 1.000 rpm, with Trypsin and LysC at a enzyme to ptotein w/w ratio of 1:50. Next day, Trypsin and LysC were added again at a enzyme to protein w/w ratio of 1:50 and proteins were digested further for 4 h at 37 °C, 1.000 rpm. Resulting peptides were acidified with 1% TFA 99% Isopropanol in a 1:1 ratio and vortexed, followed by centrifugation at 22.000 xg RT to pellet residual particles. The supernatant was transferred into a fresh tube and subjected to stage-tip clean-up via SDB-RPS. 20 μg of peptides were loaded on two 14-gauge stage-tip plugs. Peptides were washed twice with 200 μL 1% TFA 99% ddH_2_O followed by 200 μL 1% TFA 99% Isopropanol in an in-house-made Stage-tip centrifuge at 2000 xg. Peptides were eluted with 100 μL of 5% Ammonia, 80% ACN into PCR tubes and dried at 45 °C in a SpeedVac centrifuge (Eppendorf, Concentrator plus). Peptides were resuspended in 0.1% TFA, 2% ACN, 97.9% ddH_2_O. After evaluation of protein extraction efficiency, all sample preparation for Fresh, PFA-fixed, uDISCO-, 3DISCO-, Shanel-cleared tissue was performed following the SDS-SDC protocol. For LCM sample preparation, LCM samples were caught on PCR tubes with adhesive caps and successful isolation was verified by visual inspection. 20 μl of SDS-buffer was added to each tube. The tube was closed and vortex for 30 sec, followed by centrifugation for 5 min in a table-top centrifuge to ‘catch’ the LCM sample in the protein solubilization buffer, which was confirmed afterwards by visual inspection. Sample preparation was performed as described for the SDS-SDC protocol, except for the following modifications: No shaking during cooking steps; Instead of a Branson sonicator, a Bioruptor was used for each sonication step; No Covaris CP02 was used for crushing the sample; Acetone precipitation was performed at 100 μl total volume; SDC resuspension and protein digestion was performed at a 20 μl volume.

### High-pH reversed-phase fractionation

To generate a deep library of experiment-specific precursors, peptides were fractionated at pH 10 with the spider-fractionator. 50 μg of purified peptides were separated on a 30 cm C18 column in 96 min and concatenated into 24 fractions with 2 min exit valve switches. Peptide fractions were dried in a SpeedVac and reconstituted in 2% ACN, 0.1% TFA, 97.9% ddH_2_O for LC-MS analysis.

### Liquid chromatography and mass spectrometry (LC-MS)

LC-MS was performed on an EASY nanoLC 1200 (Thermo Fisher Scientific) coupled online either to a quadrupole Orbitrap mass spectrometer (Q Exactive HFX, Thermo Fisher Scientific), or a trapped ion mobility spectrometry quadrupole time-of-flight mass spectrometer (timsTOF Pro, Bruker Daltonik GmbH, Germany) via nano-electrospray ion source (Captive spray, Bruker Daltonik GmbH). Peptides were loaded on a 50 cm in-house packed HPLC-column (75 μm inner diameter packed with 1.9 μm ReproSil-Pur C18-AQ silica beads, Dr. Maisch GmbH, Germany). Sample analytes were either separated using a linear 100min gradient from 5-30% B in 80 min followed by an increase to 60% for 4 min, and by a 4 min wash at 95%, a decrease to 5% B for 4 min, and a re-equilibration step at 5% B for 4 min, or separated on a linear 120 min gradient from 5-30% B in 90 min followed by an increase to 60% for 10 min, and by a 5 min wash at 95%, a decrease to 5% B for 5 min, and a re-equilibration step at 5% B for 5 min (Buffer A: 0.1% Formic Acid, 99.9% ddH_2_O; Buffer B: 0.1% Formic Acid, 80% ACN, 19.9% ddH_2_O). Peptides derived from laser capture microdissection and matching libraries were separated using a linear 70 min gradient from 3-30% B in 45 min followed by an increase to 60% for 5 min, an increase to 95% in 5min, followed by 5 min at 95% B, a decrease to 5% B for 5 min, and an equilibration step at 5% B for 5 min. Flow-rates were constant at 300 nl/ min. The column temperature was kept at 60 °C by an in-house manufactured oven.

Mass spectrometry analysis for the evaluation of sample preparation on a Q Exactive HFX was performed in data dependent scan mode. For full proteome measurements, MS1 spectra were acquired at 60.000 resolution and a m/z range of 300-1.650 with an automatic gain control (AGC) target of 3E6 ions and a maximum injection time of 20 ms. The top 15 most intense ions with a charge of two to eight from each MS1 scan were isolated with a width of 1.4 m/z, followed by higher-energy collisional dissociation (HCD) with a normalized collision energy of 27 % and a scan range of 200-2000 m/z. MS/MS spectra were acquired at 15,000 resolution with an AGC target of 1E5, a minimum AGC target of 2.9E3, and a maximum injection time of 28 ms. Dynamic exclusion of precursors was set to 30 s.

Deep proteomes and comparisons of clearing conditions with the SDS-SDC protocol were acquired on a standard timsTOF Pro in a data-dependent PASEF mode with 1 MS1 survey TIMS-MS and 10 PASEF MS/MS scans per acquisition cycle. Ion accumulation and ramp time in the dual TIMS analyzer was set to 100 ms each and we analyzed the ion mobility range from 1/K0 = 1.6 Vs cm^−2^ to 0.6 Vs cm^−2^. Precursor ions for MS/MS analysis were isolated with a 2 Th window for m/z < 700 and 3 Th for m/z >700 in a total m/z range of 100-1.700 by synchronizing quadrupole switching events with the precursor elution profile from the TIMS device. The collision energy was lowered linearly as a function of increasing mobility starting from 59 eV at 1/K0 = 1.6 VS cm^−2^ to 20 eV at 1/ K0 = 0.6 Vs cm^−2^. Singly charged precursor ions were excluded with a polygon filter (otof control, Bruker Daltonik GmbH). Precursors for MS/MS were picked at an intensity threshold of 2.500 a.u. and re-sequenced until reaching a ‘target value’ of 20.000 a.u taking into account a dynamic exclusion of 40 sec elution.

Peptides derived from LCM samples were acquired on a timsTOF Pro modified for highest ion transmission and sensitivity, as described in Brunner et al., in a data-dependent PASEF mode with 1 MS1 survey TIMS-MS and 5 PASEF MS/MS scans per acquisition cycle. Ion accumulation and ramp time in the dual TIMS analyzer was set to 50 ms each and we analyzed the ion mobility range from 1/K0 = 1.6 Vs cm^−2^ to 0.6 Vs cm^−2^. Precursor ions for MS/MS analysis were isolated with a 2 Th window for m/z < 700 and 3 Th for m/z >700 in a total m/z range of 100-1.700 by synchronizing quadrupole switching events with the precursor elution profile from the TIMS device. The collision energy was lowered linearly as a function of increasing mobility starting from 59 eV at 1/K0 = 1.6 VS cm^−2^ to 20 eV at 1/K0 = 0.6 Vs cm^−2^. Singly charged precursor ions were excluded with a polygon filter (otof control, Bruker Daltonik GmbH). Precursors for MS/MS were picked at an intensity threshold of 1.500 a.u. and re-sequenced until reaching a ‘target value’ of 20.000 a.u taking into account a dynamic exclusion of 40 sec elution.

### Data processing

Raw files were either searched against the mouse Uniprot databases (UP00000589_10090. fa, UP00000589_10090_additional.fa) or human Uniprot databases (UP000005640_9606.fa, UP000005640_9606_additional.fa using the MaxQuant version 1.6.7.0 which extracts features from four-dimensional isotope patterns and associated MS/MS spectra. False-discovery rates were controlled at 1% both on peptide spectral match (PSM) and protein level. Peptides with a minimum length of seven amino acids were considered for the search including N-terminal acetylation and methionine oxidation as variable modifications and cysteine carbamidomethylation as fixed modification, while limiting the maximum peptide mass to 4.600 Da. Enzyme specificity was set to trypsin cleaving c-terminal to arginine and lysine. A maximum of two missed cleavages were allowed. Maximum precursor tolerance in the first search and fragment ion mass tolerance were searched as default for TIMS-DDA data. Main search tolerance was set to 20 ppm. The median absolute mass deviation for the data set was 1.57 ppm. Peptide identifications by MS/MS were transferred by matching four-dimensional isotope patterns between the runs with a 0.7 min retention-time match window and a 0.05 1/K0 ion mobility window. Label-free quantification was performed with the MaxLFQ algorithm and a minimum ratio count of 1.

### Proteomics downstream data analysis

Proteomics data analysis was performed in the Perseus environment (version 1.6.7.0), Prism (GraphPad Software, version 8.2.1). MaxQuant output tables were filtered for ‘Reverse’, ‘Only identified by site modification’, and ‘Potential contaminants’ before further processing. Protein and peptide identifications were reported after filtering as described above. Proteome correlations across technical/analytical/biological replicates were performed after log_10_-transformation. Coefficients of variation (CVs) were calculated across the full data set or within experimental groups on raw intensity levels for shared observations of more than one. Hierarchical clustering was performed in Perseus with default parameters and Pearson correlation as distance parameters. Before differential expression analysis, data were filtered for at least two observations in one group to be compared, followed by log_2_-transformation and imputation from a normal distribution modelled as the data set with a downshift of 1.8 standard deviations and a width of 0.3 standard deviations. Deep proteomes of biological replicates from fresh or vDISCO cleared tissue were tested for differences by a two-sided t-test. False-discovery rate control due to multiple hypothesis testing was performed by a permutation-based model and SAM-statistic with an S0-parameter of 0.2 and an FDR of 0.01. Ontologies for cellular compartment assignment and keywords was performed with the mainAnnot. Mus_musculus.txt.gz followed by log_2_-fold difference frequency counts for the terms ‘Extracellular space’, ‘Blood microparticle’, ‘Neurodegeneration’, ‘Aging’, ‘Neurogenesis’, ‘Receptor’, ‘Virus-Host’, ‘Immunity’, Wound healing’ and ‘Cell migration’. 1D enrichment analysis was performed on the two-sided t-test difference and only enriched terms with a size of larger than ten were displayed in the comparison of fresh versus vDISCO deep proteomes. CVs rank plots were calculated within each of the deep proteome groups and plotted against the median abundance of each protein within each group after log_10_-transformation.

For the calculation of systematic ontology-related protein mass shifts, total protein copy number estimations of the deep fresh and vDISCO cleared proteomes of biological replicates were calculated using the Perseus plugin ‘Proteomic ruler’. Protein copy numbers were calculated with the following settings: Averaging mode. ‘All columns separately’, Molecular masses: ‘Molecular weight [kDa]’, Scaling mode: ‘Histone proteomic ruler’, Ploidy: ‘2’, Total cellular protein concentration: ‘200g/l’. Proteins were annotated with regards to their cellular compartment by gene ontology from the mainAnnot. mus_musculus.txt.gz. For protein mass estimates, we multiplied the resulting protein copy number by its protein mass for each conditional replicate and summed up all protein masses to obtain the total protein mass for each representative proteome reflecting 100% of the protein mass. To calculate the subcellular protein mass contribution, we calculated the protein mass proportion for the GOCC terms related to the cytoskeleton: ‘Actin filament’, ‘Intermediate filament’, ‘Centrosome’, ‘Microtubule’; Membranes: ‘Cytoplasm’, ‘Plasma membrane’, ‘Membrane’; Organelles: ‘Mitochondrion’, ‘Nucleus’, ‘Endoplasmic reticulum’, ‘Golgi apparatus’. For calculating the organellar change between the respective Fresh and vDISCO sub-proteomes, individual protein mass contributions were normalized by its total proteome mass first, followed by ratio calculation to obtain the percentage shift of protein mass between Fresh and vDISCO brains.

For principal component analysis (PCA) of both LCM applications (mTBI and FAD), data were grouped according to their condition, filtered for at least 760 or 900 proteins for the FAD or mTBI experiment respectively and at least 2 observations within one of the two conditions, column-wise median normalized, and missing values were imputed from a normal distribution with a width of 0.3 standard deviations that was downshifted by 1.8 standard deviations. Differential expression analysis for the FAD and mTBI experiment was performed by two-sided Welch’s t-test on LFQ or IBAQ data respectively. False-discovery rate control due to multiple hypothesis testing was performed by a permutation-based model and SAM-statistic with an S0-parameter of 0 or 0.2 and FDR of 0.3 or 0.5 for the mTBI and FAD comparison, respectively.

## Movie legend

Movie S1

Step-by-step DISCO-MS pipeline illustrated on a 5×FAD mouse brain (6 weeks old).

---

**Fig. S1:**
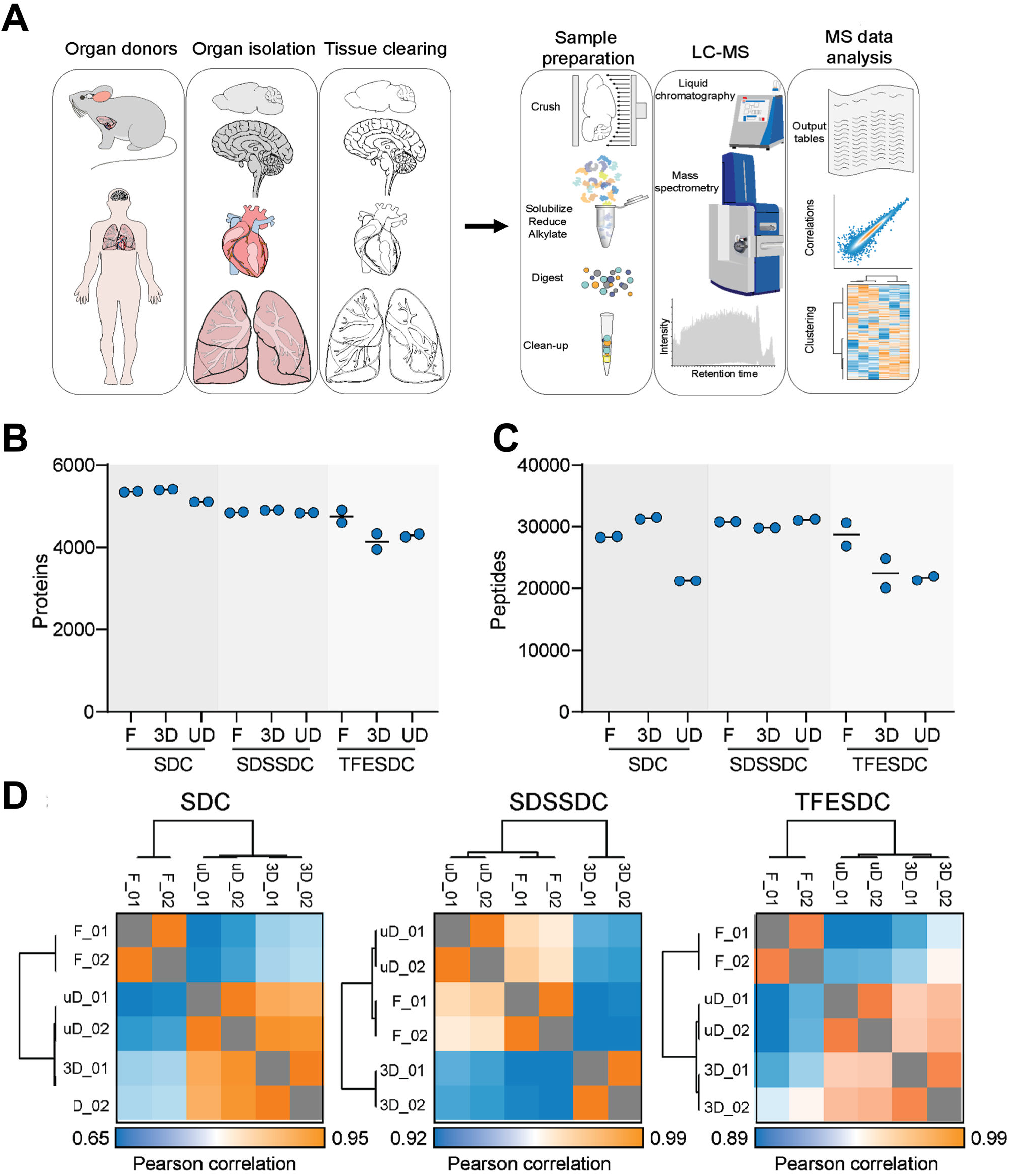
Optimization of sample preparation from cleared tissues. **(A)** The workflow for optimization of DISCO-MS from clearing bulk tissue to mass spectrometry. Organs can be isolated from any organism followed and cleared by any organic solvent-based tissue clearing. The cleared tissues then subjected to sample preparation workflow we developed for the mass spectrometry analysis. In short: the tissues were solubilized, reduced and alkylated, digested into tryptic proteins and cleaned up ready for liquid chromatography coupled to mass spectrometry (LC-MS) analysis. **(B)** Protein identifications across analytical duplicates for SDC, SDSSDC, or TFESDC preparations coming from fresh or cleared mouse brains (3D, uD: 3DISCO and uDISCO clearing methods, respectively). **(C)** Peptide levels of the samples shown in **(B)**. **(D)** Proteome correlation matrices for measurements presented in **(B)** and **(C)**. High Pearson correlations indicate very similar proteomes across conditions in SDSSDC and TFESDC.

**Fig. S2:**
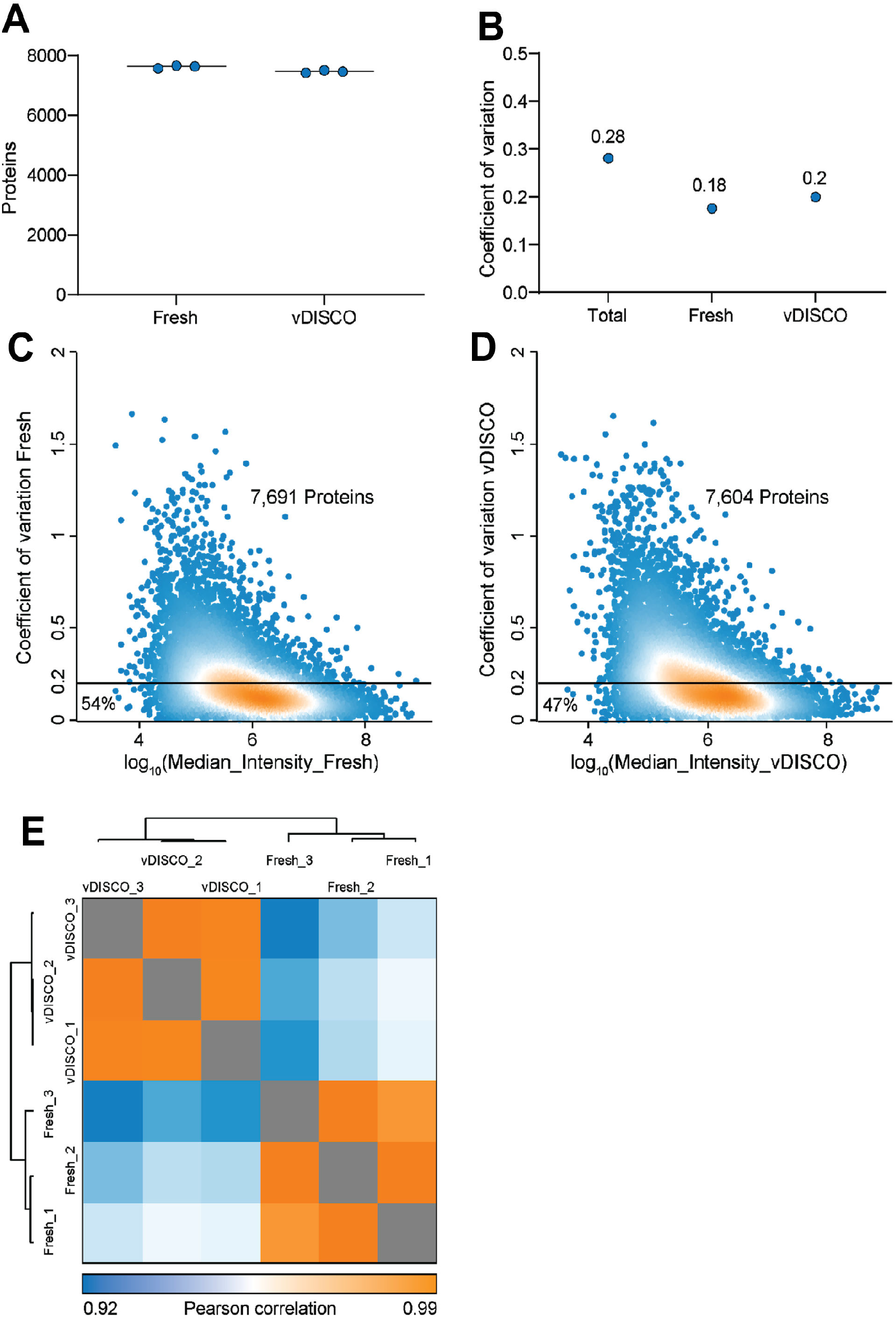
Quantitative assessment of proteome in vDISCO-cleared and fresh mouse brain tissues in biological triplicates. **(A)** Proteins identified across all three biological replicates for either fresh or vDISCO-cleared tissue. **(B)** Coefficients of variation (CV) for either the total data set including fresh and vDISCO cleared tissue, or fresh/vDISCO only. Note that CVs across biological replicates are low and that CVs across biological triplicates are very similar for fresh and for vDISCO highlighting that proteome of vDISCO-cleared organs is highly reproducible. **(C)** Abundance to CV rank plot for either fresh tissue (left; 7,691 proteins in total; 54% of all proteins are below CV = 0.2) or **(D)** vDISCO cleared tissue (right; 7,604 proteins in total; 47% of all proteins are below CV = 0.2). **(E)** Protein intensity correlation plot for all six biological replicates (3x fresh and 3x vDISCO-cleared).

**Fig. S3:**
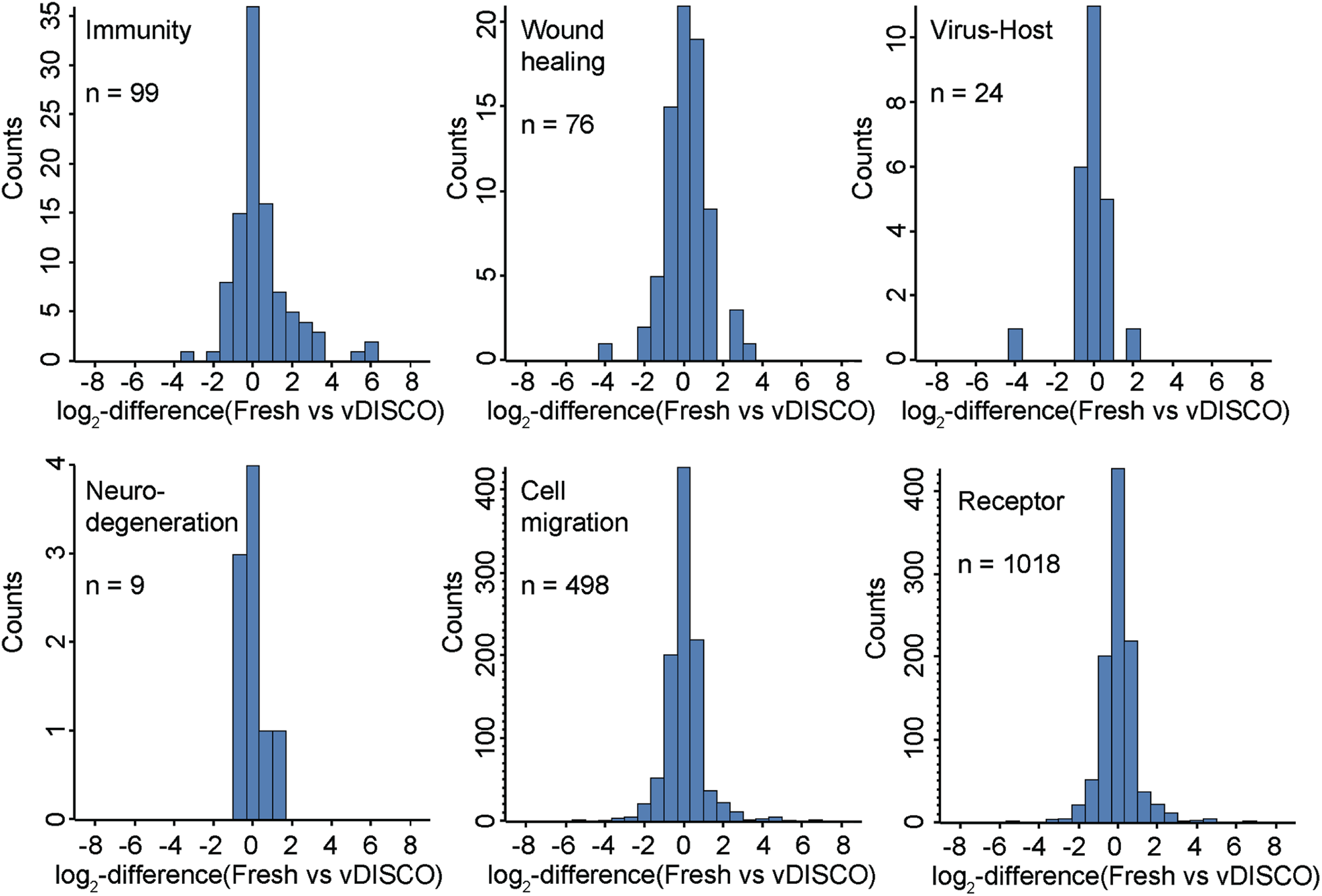
Fold-changes of gene ontologies between fresh and vDISCO cleared biological triplicate mouse brains. Log_2_-fold changes for the terms ‘Immunity’ (99 proteins), ‘Wound healing’ (76 proteins), ‘Virus-Host’ (24 proteins), ‘Neurodegeneration’ (9 proteins), ‘Cell migration’ (498 proteins) and ‘Receptor’ (1,018 proteins) between fresh and vDISCO-cleared biological triplicates.

**Fig. S4:**
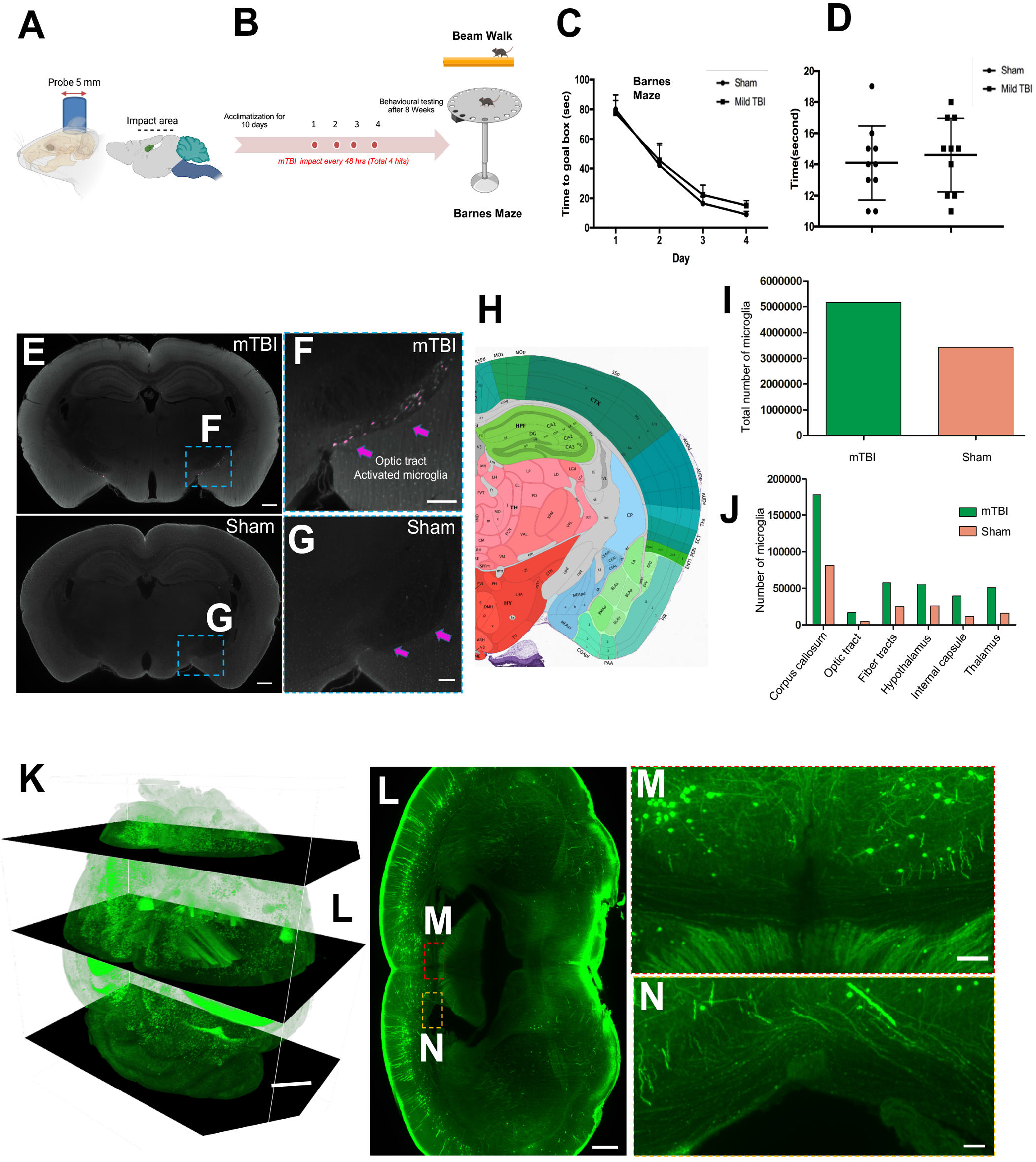
mTBI model validation by behavior and axonal morphology. **(A)** The mTBI impact area point the intact skull. **(B)** Schematic plan of repetitive mTBI experimental mouse model (red points indicate each impact time). **(C)** Barnes maze test in sham vs. mTBI animals. n=10 animals per group. **(D)** Beam walk test in sham and mTBI animals. n=10 animals per group. No significant behavior change detected-confirming the “mild” nature of our TBI model. **(E)** Coronal optical slices showing optic tract with activated microglia. Scale bar, 400 μm **(F, G)** High magnification image of optic tract in mTBI brain **(F)** vs. the same region from sham control brain **(G)** showing the activated microglia morphology in mTBI brain compared to control brain. Scale bar, 200 μm, **(H)** Corresponding brain regions (coronal view) shown in Allen Brain atlas. **(I)** Quantification of total number of microglia in mTBI vs. Sham animals. **(J)** Quantification of microglia numbers in mTBI vs. sham mice using ClearMap method. Only the regions with major changes are shown. **(K)** 3D view of whole brain from a Thy1-GFP-M after mTBI. Scale bar, 1000 μm **(L)** 2D orthoslice showing the axonal swellings in corpus callosum (white matter areas). Scale bar, 500 μm **(M, N)** High magnification images marked in **(L).** Scale bar, M, 100 μm and N, 50 μm.

**Fig. S5:**
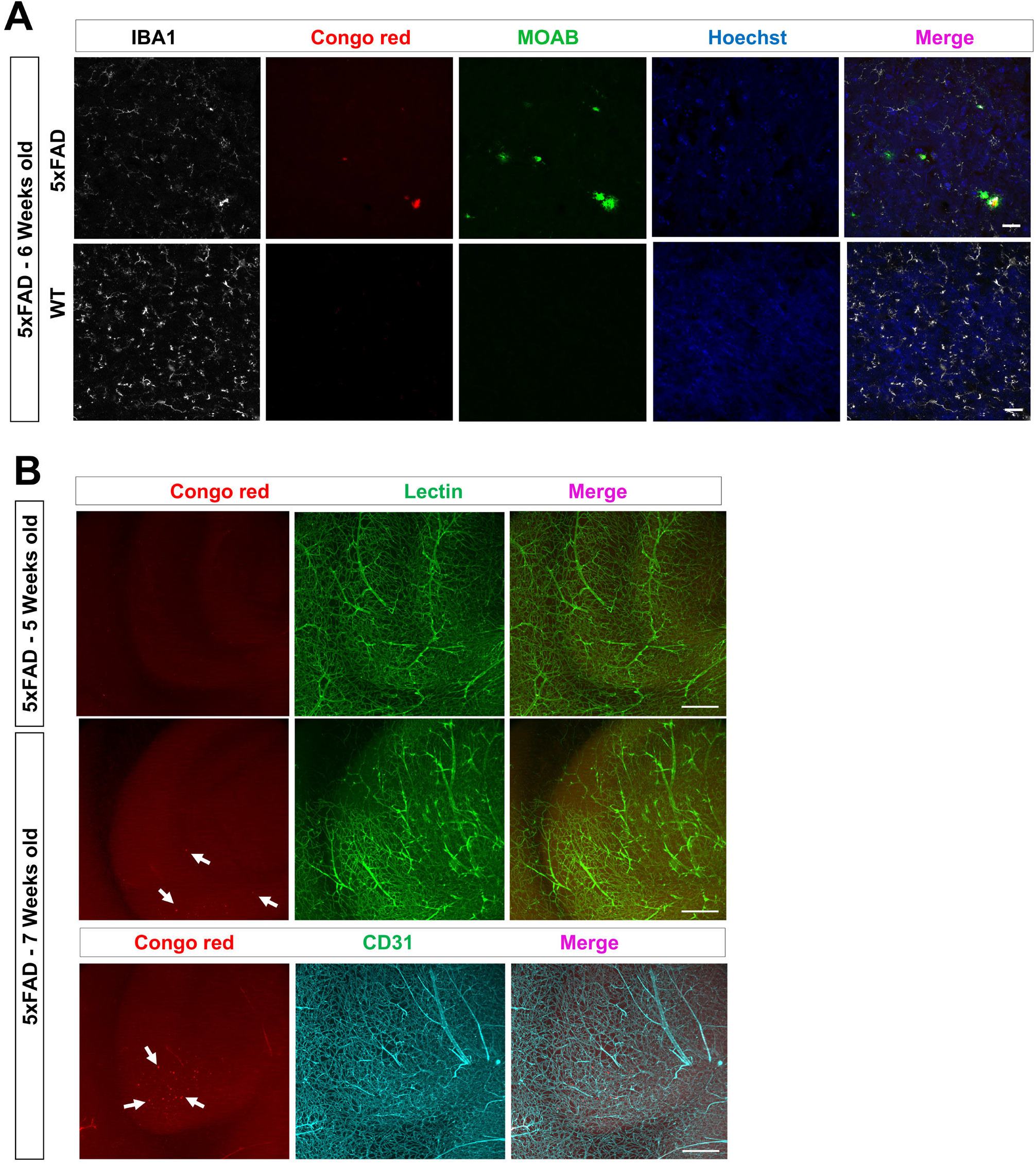
Histological validation of Aβ plaques in 5×FAD brains. **(A)** Tissue histology validation of Congo red plaque staining with a plaque-specific monoclonal antibody plaques (MOAB, green). Furthermore, the microglia were stained using IBA1 antibody (in white) and nuclei using Hoechst dye (in blue). Microglia activation around the plaques of 5×FAD mouse brain is apparent. Scale bars, 20 μm **(B)** We checked the presence of plaques in 5 weeks old 5×FAD mice by Congo red (in red) labeling along with lectin (in green) labeling of vasculature, which provides anatomical information. In contrast, we identified many plaques in 7 weeks old 5xFAD mouse brains (pointed by white arrows), whose vasculature were co-stained with lectin (in green) or CD31 antibody (in cyan). Scale bars, 200 μm.

